# Spatial heterogeneity of glioblastoma cells reveals sensitivity to NAD^+^ depletion at tumor edge

**DOI:** 10.1101/2020.11.26.399725

**Authors:** Daisuke Yamashita, Davide Botta, Hee Jin Cho, Xiaoxian Guo, Saya Ozaki, Victoria L Flanary, Inna Sirota, Mu Gao, Shinobu Yamaguchi, Mayu A Nakano, Fen Zhou, Hongyi Zhou, Toru Kondo, Takeharu Kunieda, David K Crossman, Harley I Kornblum, Myriam Gorospe, Do-Hyun Nam, Nicola Zamboni, Jeffrey Skolnick, Zhenglong Gu, Frances E Lund, Ichiro Nakano

## Abstract

Even after total resection of glioblastoma core lesions by surgery and aggressive post-surgical treatments, life-threatening tumors inevitably recur. A characteristic obstacle in effective treatment is high intratumoral heterogeneity, both longitudinally and spatially. Recurrence occurs predominantly at the brain parenchyma-tumor core interface, a region termed *tumor edge.* Given the difficulty of accessing it surgically, the composition of the tumor edge, harboring both cancerous and non-cancerous cells, remains largely unknown. Here, to identify phenotypic diversity among heterogeneous glioblastoma core and edge lesions, we uncovered the existence of three phenotypically-distinct clonal subpopulations within individual tumors from glioblastoma patients. Clones from the tumor core shared the same phenotype, exclusively generating tumor-core cells. In contrast, two distinct clonal subtypes were identified at the tumor edge: one generated only edge-lesion cells and the other expanded more broadly to establish both edge- and core-lesions. Using multiple xenograft experimental models in mouse brains, tumor edge development was found to require that both somatic and tumor cells express the NADase CD38, combinedly elevating glioblastoma malignancy. *In vitro* data suggested that intracellular NADase activity at the edge was provoked through intercellular communication between edge clones and normal astrocytes. Systemic treatment of tumor-bearing mice with 78c, a small-molecule CD38 inhibitor, attenuated the formation of glioblastoma edge lesions, suggesting its clinical potential to pharmacologically eliminate tumor-edge lesions. Collectively, these findings provide novel phenotypic and mechanistic insights into clonal heterogeneity within glioblastoma, particularly in the surgically unresectable, currently understudied tumor edge.

## INTRODUCTION

Surgical resection is the first-line treatment for glioblastoma, the most common and lethal intraparenchymal cancer in adult patients. Nonetheless, the infiltrative nature of tumor cells makes their complete removal an impractical goal. Recent studies suggested that early-branched ancestor-like tumor cells were specifically located at tumor edge, which may act as progenitor cells for tumor core-located cells in single glioblastoma tumors^1,2^. Likewise, data in multiple murine glioblastoma-like tumor models has supported this Edge-to-Core (E-to-C) progression theory^3^. In addition, the clinical significance of the E-to-C molecular axis was indicated in the primary-to-recurrent glioblastoma setting with our cohort^4^. However, whether this theory universally explains the development of primary glioblastoma remains unknown. Because multi-OMICs analyses have identified largely distinct signaling mechanisms activated in edge- and core-located tumor cells (e.g. Esm1/endocan, Bruton’s Tyrosine Kinase, Nitrogen metabolism)^3–9^, establishing spatially distinct therapeutic modalities to concur glioblastoma is a critical challenge.

The spatial identity of individual tumor cells programmed to create tumor-edge and tumor-core structures has just started to be elucidated by newly established models of tumor edge and core from both patient-derived and murine tumors^3–5^. Edge-located brain tumor cells exist in a highly specified ecosystem, quite distinct from that of the tumor core, harboring active interaction with various somatic cells such as neurons^10–12^, astrocytes^13^, vascular endothelial cells^3^, and immune cells^14^. These tumor-associated somatic cells may contain cellular populations that contribute to activating or suppressing tumor cells. Yet, unlike the murine tumor system^3^, it remains unknown whether signals from somatic cells in the tumor microenvironment help shape the edge lesions of tumors. Such intercellular signals at the tumor edge could constitute therapeutic targets. Molecular investigation is required to understand the phenotypic dynamics of patient tumor cells.

The glycoprotein CD38 is located in distinct subcellular regions including mitochondria. Mitochondrial CD38 reduces the levels of nicotinamide adenine dinucleotide (NAD^+^), an essential metabolite for energy homeostasis and DNA repair^15^ in both somatic cells and cancer cells. Nicotinamide mononucleotide (NMN) is a plasma metabolite in the mammalian NAD^+^ biosynthesis pathway, and a key intermediate synthesized from nicotinamide, 5’-phosphoribosyl-1-pyrophosphate or nucleoside riboside^16,17^ to effectively elevate NAD^+^ biosynthesis in mitochondria^18^. Re-activation of the NAD^+^ pathway in the aging brain by supplementation of NMN appears to alter the microenvironment leading to resistance, at least partially, to tumor-burden death^8^. These data raise the possibility that the titration of NAD^+^ activity *via* CD38 in tumor cells and adjacent somatic cells is associated with glioma malignancy.

In this study, we sought to define the cellular and molecular heterogeneity in the glioblastoma-edge microenvironment. We further sought insight into therapeutic interventions directed at the key molecules driving edge formation in glioblastoma. To do so, we first isolated multiple tumor clones from patient glioblastoma edge and core tissues and characterized them phenotypically, particularly in the context of intratumoral spatial identity.

## RESULTS

### Distinct spatial identities of glioblastoma cells at the core and tumor edge

First, we investigated whether patient tumor cells at the edge and core retain their spatial identity when placed into mouse brains as xenografts. T 1-Gadlinium-enhancing tumor core and non-enhancing T2/FLAIR-high subcortical edge tissues (ptCore and ptEdge, respectively) were collected through supra-total resection in awake surgery for glioblastoma patients using MRI-guided biopsy, followed by histopathological confirmation (**Fig. 1a and Extended Data Fig. 1a, b**)^5,6,9^ In particular, ptEdge were secured from the deeper subcortical area of the brain, one of the most frequent brain regions to cause post-craniotomy tumor recurrence^4,19^. ptCore- and ptEdge-derived single cells were then labeled with lentiviral vectors to express green fluorescence protein (GFP) or mCherry, respectively (**Fig. 1b**). Immediately after lentiviral infection, these tumor cells were injected into mouse brains (i.e. GFP-expressing patient core cells and mCherry-expressing patient edge cells) (**Fig. 1b and Extended Data Fig. 1c**). Although multiple cases (n>5) were used to attempt this approach, subsequently only one glioblastoma case (g1051) resulted in lethal tumor formation in mouse brains. As early as at day 2 after injection, xenografted cells have already started to create core and edge lesions (msCore and msEdge, respectively), which generated larger tumors by day 30 (**Fig. 1c, d**). Tumor immunohistochemistry (IHC) with anti-GFP and anti-mCherry revealed that msEdge lesions are predominantly formed by mCherry^+^ ptEdge cells, while msCore lesions contained both GFP^+^ ptCore cells and mCherry^+^ ptEdge cells at both time points (Fig. 1e, f). In addition, ptEdge cells were preferentially located in two specific brain regions: in the corpus callosum and at the subventricular zone (SVZ) adjacent to the lateral ventricle, two major brain regions of glioblastoma infiltration in patients (**Extended Data Fig. 1d, e**). Quantification of dissociated tissues from msCore and subcortical msEdge by flow cytometry for GFP and mCherry revealed that msEdge harbored both ptEdge-derived cells (5.86%) and ptCore-derived cells (7.49%), while msEdge mostly comprised ptEdge-derived cells (17.2% *vs.* 1.47%) (**Fig. 1g**). Despite the limitation of the case number, these introductory results are consistent with the IHC data and suggest that, similar to the murine tumor models^3^, ptEdge cells comprise two distinct cellular populations with the capacity to generate tumor core and tumor edge lesions, while the majority of ptCore cells are programmed to form only the core lesions.

**Fig. 1:**
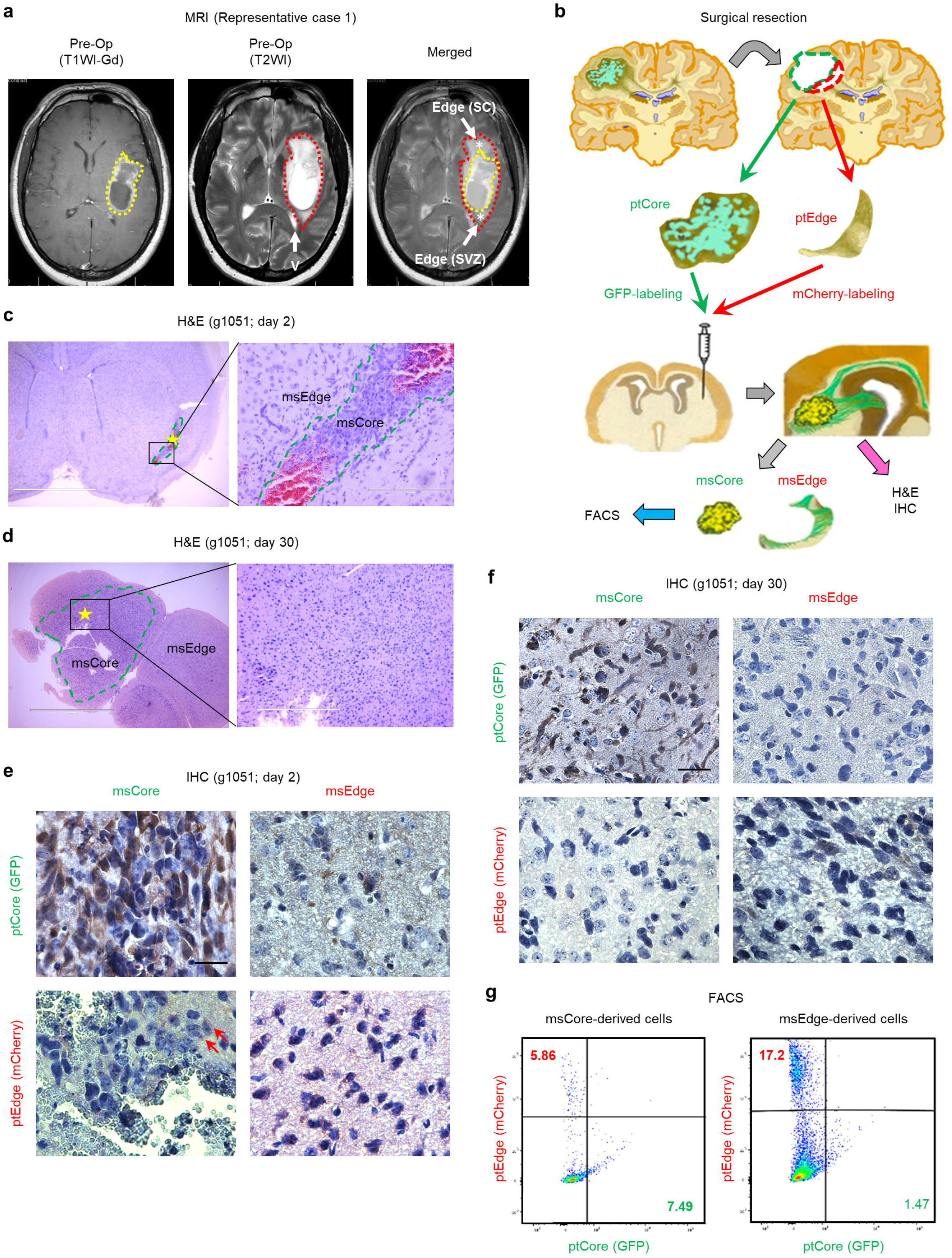
Distinct spatial identities of the tumor edge and core cells in glioblastoma. (**a**) Representative Magnetic Resonance Imaging (MRI) of a glioblastoma patient depicting the glioblastoma core area (yellow) and edge areas (red). V: lateral ventricle; SC: subcortical edge; SVZ: subventricular zone. (**b**) Schematic for surgical resection of ptCore and ptEdge, tumor challenge in mouse brains after lentiviral labeling, and subsequent analyses. (**c**) Representative H&E staining of mouse brains harboring g1051 ptCore and ptEdge tumor cells 2 days after intracranial injection. Scale bars represent 2 mm (left) and 200 μm (right). (**d**) Representative H&E staining of mouse brains harboring g1051 ptCore and ptEdge tumor cells 30 days after intracranial injection. Scale bars represent 2 mm (left) and 200 μm (right). (**e**) Representative IHC imaging for GFP (top) and mCherry (bottom) in msCore (left) and msEdge (right) 2 days after intracranial injection. Scale bar represents 100 μm. (**f**) Representative IHC imaging for GFP (top) and mCherry (bottom) in msCore (left) and msEdge (right) 30 days after intracranial injection. Scale bar represents 100 μm. (**g**) Fluorescence activated cell sorting (FACS) for GFP (indicvative for ptEdge cells) and mCherry (indicative for ptCore cells) using msCore-derived cells (top) and msEdge-derived (bottom) tumor cells.

### Individual glioblastoma tumors comprise three clonal populations harboring distinct spatial identities

Next, we investigated whether the spatial identities of the three different tumor cell populations (ptCore, core-located ptEdge, and edge-located ptEdge) are in fact a consequence of the presence of three distinct clonal subpopulations. Among more than ten cases attempted, we succeeded in establishing multiple clonal lines from three patients (g1051, g101027, g303016) by using the serial dilution method individually from the matched ptCore and ptEdge samples of each tumor (total established clonal n = 46) (**Fig. 2a and Extended Data Fig. 2a**). To validate that these cultures were actually single cell-derived clones, we performed DNA sequencing to detect single-nucleotide polymorphisms (SNP) within each clone. This analysis confirmed that each of the established clones (tested n=15) had a unique SNP signature and was a *bona fide* individual clone (**Extended Data Fig. 2b**). To uncover the phenotypic differences among these clones, ptCore (n=7) and ptEdge (n=6) clones were injected into the brains of immunodeficient mice to follow their spatial identities. In the ptCore group, IHC for human mitochondria revealed that they formed a core-like tumor structure at the injection site (msCore) (**Fig. 2b**), akin to the results seen after direct injection (**Fig. 1**). In contrast, ptEdge clones exhibited two distinct populations: one migrating subcortically from the injection site primarily toward the corpus callosum and lateral ventricle without core formation (msEdge), and the other generating both msCore and msEdge, indicating their transitional (or bi-functional) phenotype (termed trans-ptEdge) from Edge to Core or *vice versa* (**Fig. 2c and Extended Data Fig. 2c**). Because vascular density is a major microenvironment-composing factor between the tumor edge and the core^3^ we next employed the slice culture assay with neonatal mouse brains to define the proximity of these glioblastoma clones with vasculature (**Fig. 2d**). When incubated with brain slices, ptEdge clones were tightly associated with the brain vasculature, whereas both ptCore and trans-ptEdge clones were randomly distributed across the brain slices without noticeable affinity for vasculature (**Fig. 2c, d**).

**Fig. 2:**
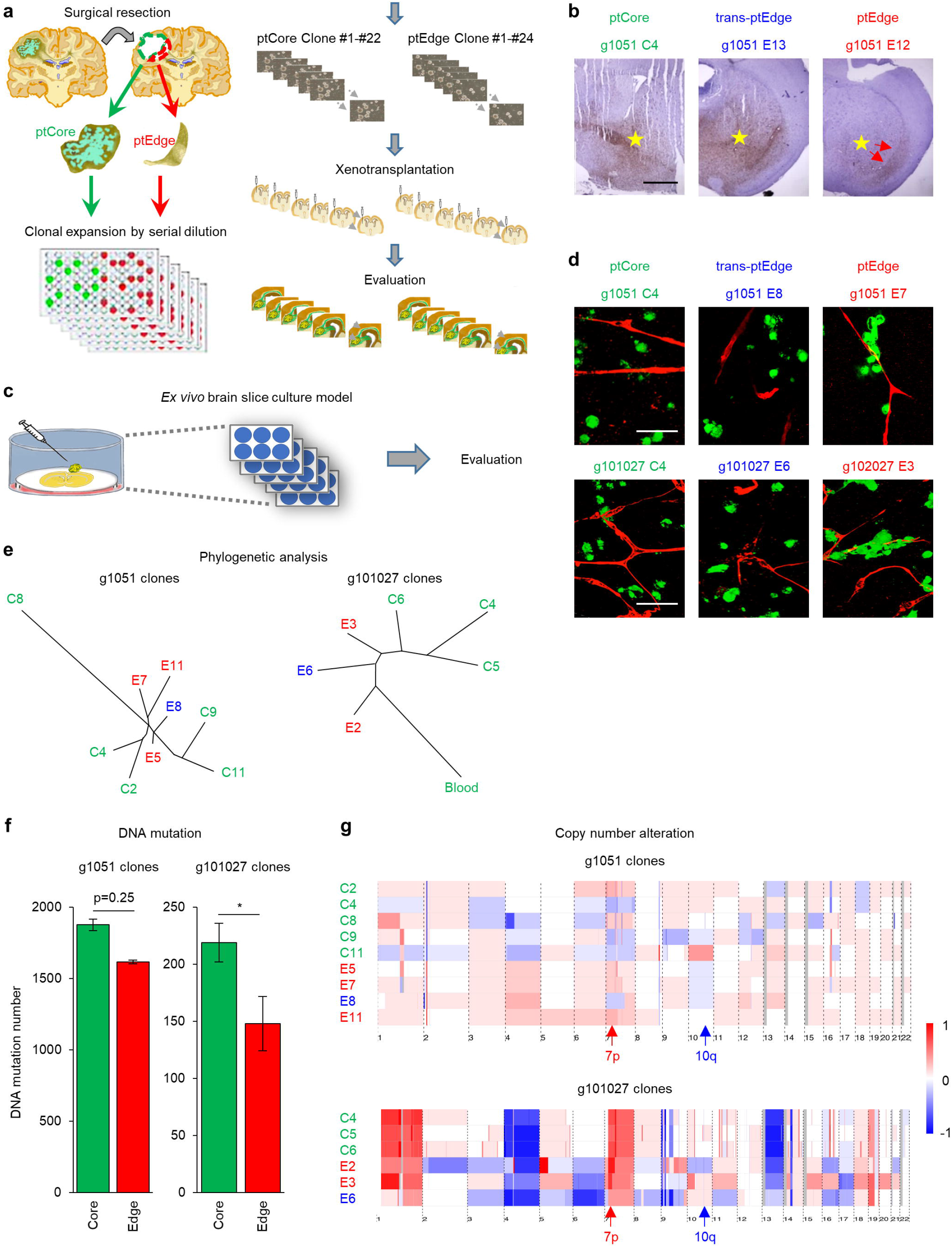
Individual glioblastoma tumors are composed with three clonal populations harboring distinct spatial identities. (**a**) Schematic demonstrating the experimental flow to establish ptCore and ptEdge clones. (**b**) Representative IHC for human mitochondria to label tumors derived from g1051 Core (left), trans-Edge (middle), and Edge (right) clones. Injection site indicated in yellow asterisks. Red arros indicate the infiltrating tumor cells within corpus callosum. Scale bar represents 1 mm. (**c**) Schematic demonstrating the experimental flow for *ex vivo* brain slice cultures using two ptCore and ptEdge clones. (**d**) Representative immunofluorescence with slice cultures of mouse brains together with g1051 Core (left), trans-Edge (middle), and Edge (right) clones (green). Vessels were stained with collagen IV antibody (red). Scale bar represents 20 μm. (**e**) Phylogenetic analysis of Core (green), trans-Edge (blue), and Edge (red) clones from g1051 (left) and g101027 (right) based on shared clone-specific and somatic mutations (blood). (**f**) Graphical representation of DNA mutation burdens in g1051 (left) and g101027 (right) Core and Edge clones (n=9 and 6, respectively). (**g**) Heatmap analysis comparing copy number alterations in individual chromosomes with g1051 (top) and g101027 (bottom) clones.

Given the identification of three phenotypically distinct clonal populations in glioblastoma, we then examined whether the core clones give rise to their edge counterparts or *vice versa.* Phylogenetic analyses based on DNA mutation data from individual clones demonstrated that the ptEdge clones were more genetically similar and ancestor-like, whereas the ptCore clones were more genetically diverse and progeny-like (**Fig. 2e and Extended Data Fig. 2d**). Consistent with these data, the accumulation of DNA mutations was higher in the ptCore clones than the ptEdge, among which largely overlapping mutations were observed to varying degrees between two patient-derived models (**Fig. 2f and Extended Data Fig. 2e**). A comparison of single-gene mutations demonstrated that generalized cell line mutations in g1051 were present in commonly mutated genes such as *BRCA2,* although this trend was less obvious in g101027 (**Extended Data Fig. 2f**). Based on hierarchical clustering to compare the copy number alterations of mutated genes in these clones, the most frequently dysregulated chromosomes in glioblastoma, chromosomes 7p and 10q, exhibited the highest diversity in patterns of mutations (**Fig. 2g and Extended Data Fig. 2g**). Collectively, these data suggest that the ptEdge clones diverge early in tumor development, originating trans-ptEdge clones as a reservoir for ptCore clones, collectively elevating clonal heterogeneity through tumor evolution.

### Mitochondria function is dysregulated only in ptEdge clones

Given the cell-autonomous phenotypic differences of these clonal lines, we investigated their molecular profiles to identify the mechanisms that govern intracellular spatial identity, specifically by RNA-sequencing (RNA-seq) and metabolite mass spectroscopy. For RNA-seq, we used 12 ptCore, 4 trans-ptEdge, and 5 ptEdge lines in addition to widely utilized conventional glioma spheres obtained from our institute (n=22 in total). As reference, we also recovered 3 pairs of glioblastoma tissue samples of tumor core and subcortical edge at surgery, as well as non-neoplastic cortical tissue samples from epilepsy surgery from 4 other patients (n=10 in total) (**Fig. 3a**). Principal component analysis (PCA) of tissue-derived RNA-seq data revealed a gene signature axis of “Normal tissues-Edge tissues-Core tissues” (**Fig. 3a and Extended Data Fig. 3a**). In comparison to these tissue data, the gene signatures of both ptCore clones and conventional glioma spheres were relatively close to Core tissues – indicating that these models define the characteristics of tumor cells located within the core (**Fig. 3a**). In contrast, the gene signatures of ptEdge clones were highly heterogeneous and substantially apart from any other samples (**Fig. 3a and Extended Data Fig. 3b, c**). Interestingly, the gene signatures of trans-ptEdge clones appeared to localize between those of ptEdge and ptCore clones (**Fig. 3b**), indicating that the transcriptomic features of these three types of clones reflect their intratumoral spatial identity. Gene set enrichment analysis (GSEA), Gene ontology (GO), and Ingenuity Pathway Analysis (IPA) revealed a shared activation of the *Myc* and *Hypoxia/HIF1α* pathways and a variety of metabolic pathways in ptCore tissue samples and clones (**Fig. 3c and Extended Data Fig. 3d-i**). In contrast, for ptEdge there was less overlap in pathways between the tissues and clones, presumably due to the large difference in purity of tumor cells between these two sets of samples (5-10% *vs.* 100%). Among those that were identified, mitochondrial Ca^+^ signaling pathways and extracellular matrix-associated pathways were represented in ptEdge tissue and clones, respectively (**Fig. 3c and Extended Data Fig. 3f**).

**Fig. 3:**
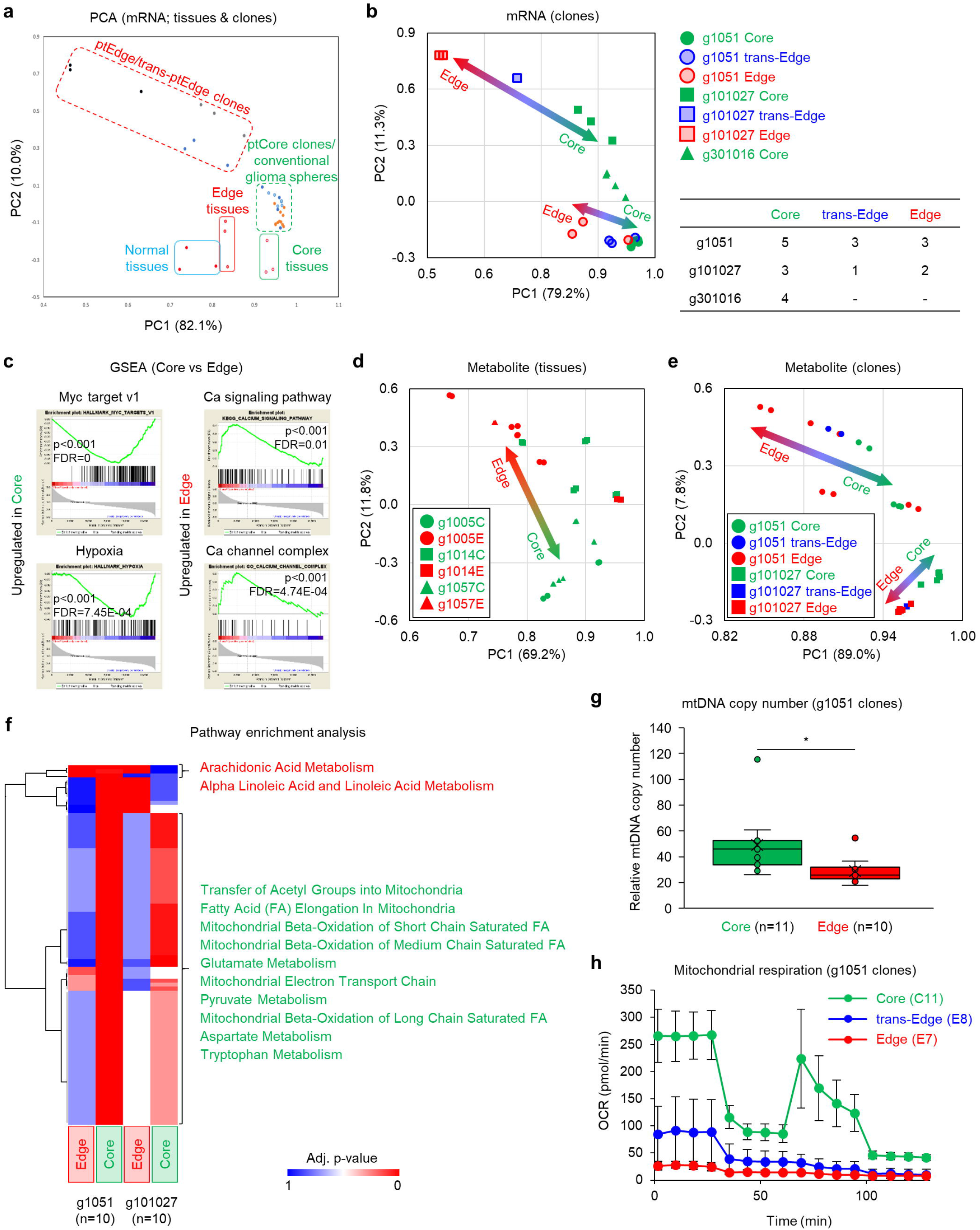
Mitochondria is dysregulated in ptEdge clones, unlike ptCore ones. (**a**) Principal component analysis (PCA) of RNA-seq data comparing normal brain tissues (n=4), matched ptCore and ptEdge tissues (n=6), 21 clones (12 Core (blue and light-blue), 4 trans-Edge, and 5 Edge), and 22 conventional glioma sphere lines (orange). (**b**) PCA of RNA-seq data comparing the indicated g1051, g101027, and g301016 clones and table for classification of each clone. (**c**) Gene set enrichment analysis (GSEA) indicating the most upregulated pathways in ptCore and ptEdge tissues. (**d**) PCA comparing metabolite expression profiles in g1005, g1014 and g1057 ptCore and ptEdge tissues (n=36 in total). (**e**) PCA comparing metabolite expression profiles in g1051 and g101027 Core, trans-Edge, and Edge clones (n=36 in total). (**f**) Metabolic pathway enrichment analysis comparing the most upregulated pathways in g1051 and g101027 Core and Edge clones (n=10 in each). (**g**) Mitochondrial DNA (mtDNA) copy number by mtDNA seq using g1051 Core and Edge clones (n=21). \**p*<0.05. (**h**) Mitochondrial respiration in g1051 Core, trans-Edge, and Edge clones, measured by oxygen consumption rate (OCR).

Given the observed differences between ptCore and ptEdge cells in metabolic pathways, we performed mass spectroscopy for metabolite profiling by using ptCore and ptEdge tissues and clones (total n=18)(**Extended Data Fig. 3j-m**). The results displayed an axis of metabolite expression between the glioblastoma edge and core (**Fig. 3d, e**), adding another layer of the molecular E-to-C axis to the RNA-seq data (**Fig. 3b, e**). Pathway analysis identified downregulation of several mitochondrial metabolites and mitochondrial pathways in ptEdge (**Fig. 3f and Extended Data Fig. 3n-p**). To support this data, mitochondrial DNA (mtDNA) levels were significantly lower in the ptEdge clones than in ptCore clones (**Fig. 3g and Extended Data Fig. 3q**). Furthermore, oxygen consumption rate (OCR), proton leak across the inner mitochondrial membrane, and ATP production were all pathologically dysregulated in ptEdge clones relative to ptCore clones (**Fig. 3h and Extended Data Fig. 3r, s**). Taken together, ptEdge cells exhibit a molecular phenotype defined by the mitochondrial dysfunction unlike ptCore cells.

### Elimination of somatic CD38 in the brain vanishes tumor edge lesions

We then sought to determine the key molecules responsible for *‘edgeness’* in glioblastoma. Among 20,787 protein-coding genes in the RNA-seq data, *CD38* represented the only gene with statistically significant expression increase in patient edge tissues (**Fig. 4a**). Volcano plot analyses validated the edge-specific elevation of *CD38* mRNA expression levels in comparison to both tumor core and normal brain tissues (**Fig. 4b and Extended Data Fig. 4a, b**). A publicly available dataset (the Ivy GBM Atlas database) verified that *CD38* mRNA expression levels were higher at the leading edge of glioblastomas (n=246) (**Fig. 4c**). Consistently, IHC of patient tumors (n=19) exhibited an enrichment of CD38 protein at tumor edge (**Fig. 4d, e**).

**Fig. 4:**
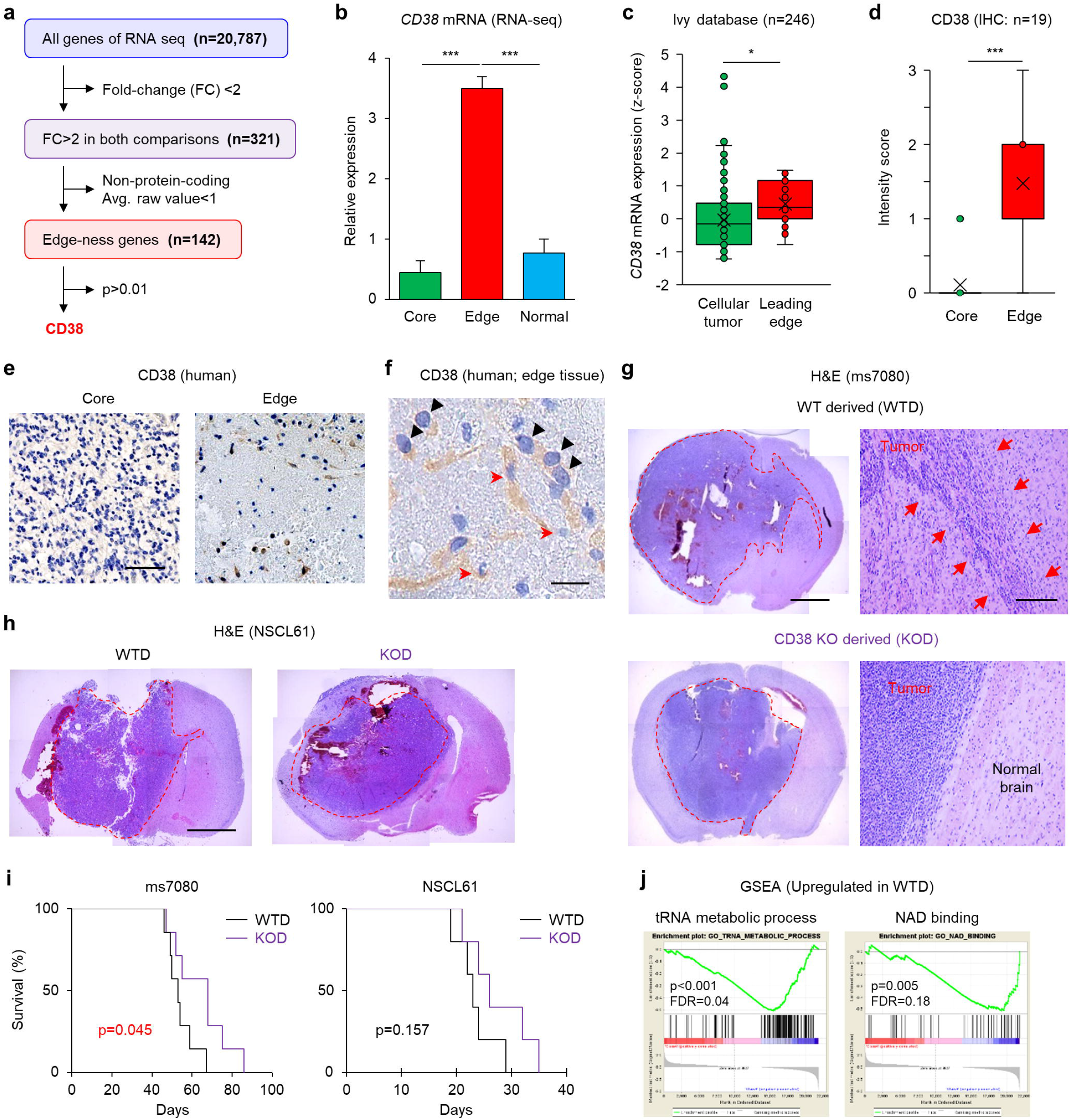
Elimination of somatic CD38 in the brain vanishes tumor edge lesions. (**a**) Schematic demonstrating the filtering procedure for 20,787 genes, with fold-change > 2 and *p* value <0.01 as markers for glioblastoma edge phenotype. *p*<0.01 for CD38 using Student’s *t-* test. (**b**) Bar graph comparing relative expression of CD38 by RNA-seq of ptCore, ptEdge, and normal brain tissues. \*\*\**p*<0.001. (**c**) Graphical comparison of CD38 expression in glioblastoma cellular tumor lesion and leading edge in the IVY Atlas database. n=246. \**p*<0.05. (**d**) Intensity score of IHC imaging for CD38 in human glioblastoma tissues. n=19. \*\*\**p*<0.001. (**e**) Representative IHC imaging for CD38 in human glioblastoma tissues. Scale bar represents 100 μm. (**f**) Representative IHC imaging for CD38 in ptEdge tissue. Edge glioblastoma cells are indicated with black arrowheads and neighboring astrocytes are indicated with red arrows. Scale bar represents 100 μm. (**g**) Representative H&E staining of mouse brains harboring WT-derived (WTD; left) and CD38KO-derived (KOD; right) ms7080 tumors. Red arrows indicate infiltrative tumor cells at the edge in WTD tumors. Scale bars represent 2 mm (left) and 200 μm (right). (**h**) Representative H&E staining of mouse brains harboring WTD (left) and KOD (right) NSCL61 tumors. Scale bars represent 2 mm (left) and 200 μm (right). (**i**) Kaplan Meier analysis comparing the survival of WTD- and KOD-tumor bearing mice (left; ms7080, n=7 in each group, right; NSCL61, n=5 in each group). (**j**) GSEA plot analysis identifying the upregulated pathways in WTD and KOD tumors.

We then sought to determine the role of CD38 in the generation of tumor edge lesions. First, we examined the cell types expressing CD38 at tumor edge *in situ.* IHC analysis in patient tumors identified ptEdge tumor cells and neighboring astrocytes as the predominant cell types expressing CD38 (**Fig. 4f**). The data from the publicly available scRNA-seq database of normal adult mouse brain (http://betsholtzlab.org/VascularSingleCells/database.html) supported this observation (**Extended Data Fig. 4c**)^20^. After confirming the NADase activity of CD38 in astrocytes from WT but not CD38 KO mice (**Extended Data Fig. 4d, e**), we investigated whether the tumor edge formation requires astrocytic and/or tumor CD38. First, CD38 KO and WT animals, harboring no noticeable phenotypic differences in brain morphology, were subjected to intracranial injection of 2 different glioma sphere models (ms7080 and NSCL61^21^). Strikingly, in both cases, glioblastoma tumors in WT mice (‘WTD tumors’) harbored tumor core lesions with extensive subcortical tumor infiltrating edge lesions, in the sharp contrast to CD38 KO mice (‘KOD tumors’) failing to generate edge lesions (**Fig. 4g, h**). Importantly, the survival of KOD tumor-bearing mice improved significantly, suggesting that impairing the tumor edge phenotype reduces the lethality of glioblastoma (**Fig. 4i**). GSEA of RNA-seq data with these tumors identified reduced NAD binding and tRNA metabolism in KOD tumors, highlighting a role for CD38 as the NADase to govern tumor edge and core heterogeneity (**Fig. 4i and Extended Data Fig. 4f**). Consistent with these data, RNA-seq analysis of non-cancerous cortical tissues from CD38 KO mice identified reductions in mitochondria-related pathways accompanied by the increases in Ca^+^ signaling pathway and arachidonic acid metabolism (**Extended Data Fig. 4g**). Collectively, these data suggest that the somatic NADase CD38 is mandated to establishing glioblastoma-edge lesions in mouse brains.

### Diminishing tumor CD38 activity attenuates tumor edge formation

Next, we investigated whether tumor-CD38 also plays a role in the tumor edge formation. Following the establishment of tumor cell cultures from WTD and KOD tumors (**Fig. 5a**), we found that KOD tumor cells exhibited substantially lower, but not absent, NADase (CD38) activity (**Fig. 5b**). Thus, we injected WTD and KOD cells harboring the distinct levels of CD38 activities into BL6 WT mouse brains. Tumors derived from KOD cells (termed CD38^low^ tumor cells) formed only partial tumor edge lesions with the core lesions harboring central necrosis, in stark contrast to the tumor architecture derived from WTD cells (termed CD38^high^ tumor cells), which displayed distinct tumor core and edge (**Fig. 5c**). As expected, mice with CD38^low^-derived tumors had significantly shorter survival than mice with CD38^high^-derived tumors (**Fig. 5d**). Collectively, whether expressed from somatic cells or tumor cells, CD38 critically promotes tumor edge formation, thereby elevating glioblastoma aggressiveness *in vivo.*

**Fig. 5:**
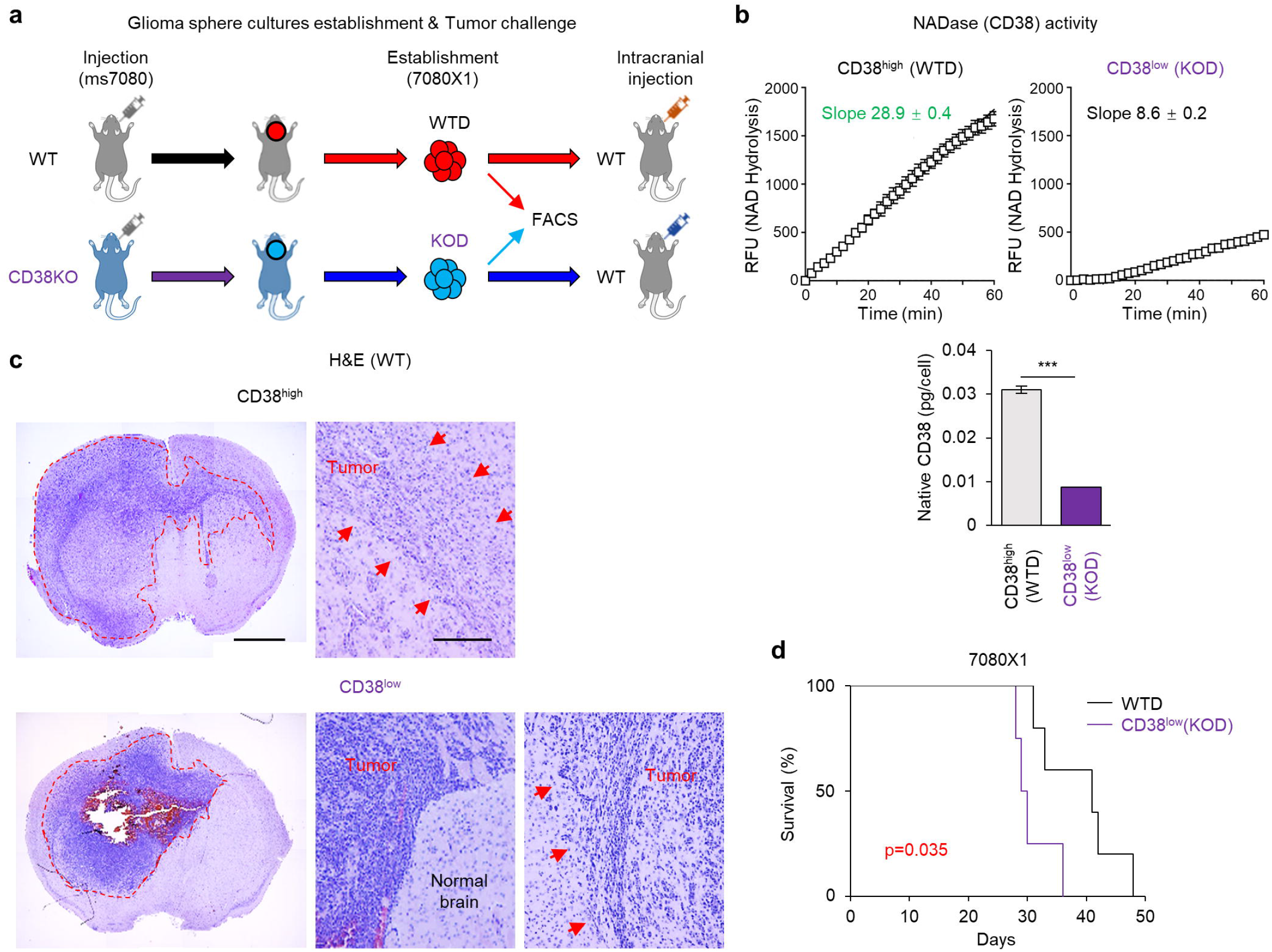
Diminished tumor-CD38 partially destroy the tumor edge formation. (**a**) Schematic of the establishment of WTD and KOD sphere cultures for intracranial injection. (**b**) Cell-surface NADase activity in WTD (left) and KOD (right) glioblastoma cells, measured by relative fluorescence units (RFUs) versus incubation time in minutes. (**c**) Representative H&E of mouse brains following intracranial injection of WTD (left) and KOD (right) glioblastoma cells. Scale bars represent 2 mm (left) and 200 μm (right). (**d**) Kaplan-Meier analysis comparing the survival of tumor-bearing mice following intracranial injection of WTD (left) and KOD (right) glioblastoma cells (n=7 for WTD glioblastoma, n=5 for KOD glioblastoma).

### Elevated CD38 activity at tumor edge cells

Distinct function of CD38 is recognized at three different regions within a cell; nucleus, cell surface, and mitochondria^22^. In order to uncover its specific function in ptEdge cells, we first performed immunofluorescence of tumor tissue, which showed that CD38 in the tumor edge tissue samples was specifically colocalized with the mitochondrial marker COX4 (**Fig. 6a**). Consistently, CD38 expression in mitochondria and nuclei by cell fractionation followed by western blot analysis revealed that mitochondrial CD38 was clearly more abundant in ptEdge clones than in ptCore clones, while nuclear CD38 was more or less similar between the two cell populations (**Fig. 6b, c and Extended Data Fig. 6a-c**). Regarding the cell surface CD38, FACS analysis showed that both patient-derived glioblastoma clones and mouse tumor models (WTD and KOD cells) yielded little or no expression of CD38 in both edge and core cells (**Extended Data Fig. 6d, e**). Given that the NADase activity is the predominant function of mitochondrial CD38, we utilized luciferase assays to detect the ratio of total oxidized and reduced NAD (NAD^+^ and NADH, respectively) indicative of the NAD activities between these clones. As expected, overall NAD activity was elevated in the ptCore clones compared to the edge counterparts (total clone n=12) (**Fig. 6d and Extended Data Fig. 6f**). Collectively, the total CD38 expression was significantly higher in the ptEdge clones compared to the ptCore and trans-ptEdge counterparts (**Fig. 6e and Extended Data Fig. 6g**). Likewise, the NADase activity on cell surface was upregulated in the tested ptEdge clones as compared to the others (**Fig. 6f and Extended Data Fig. 6h**). Taken together, these data suggest that intratumoral CD38 is primarily activated in the ptEdge cells, particularly within their mitochondria as NADase.

**Fig. 6:**
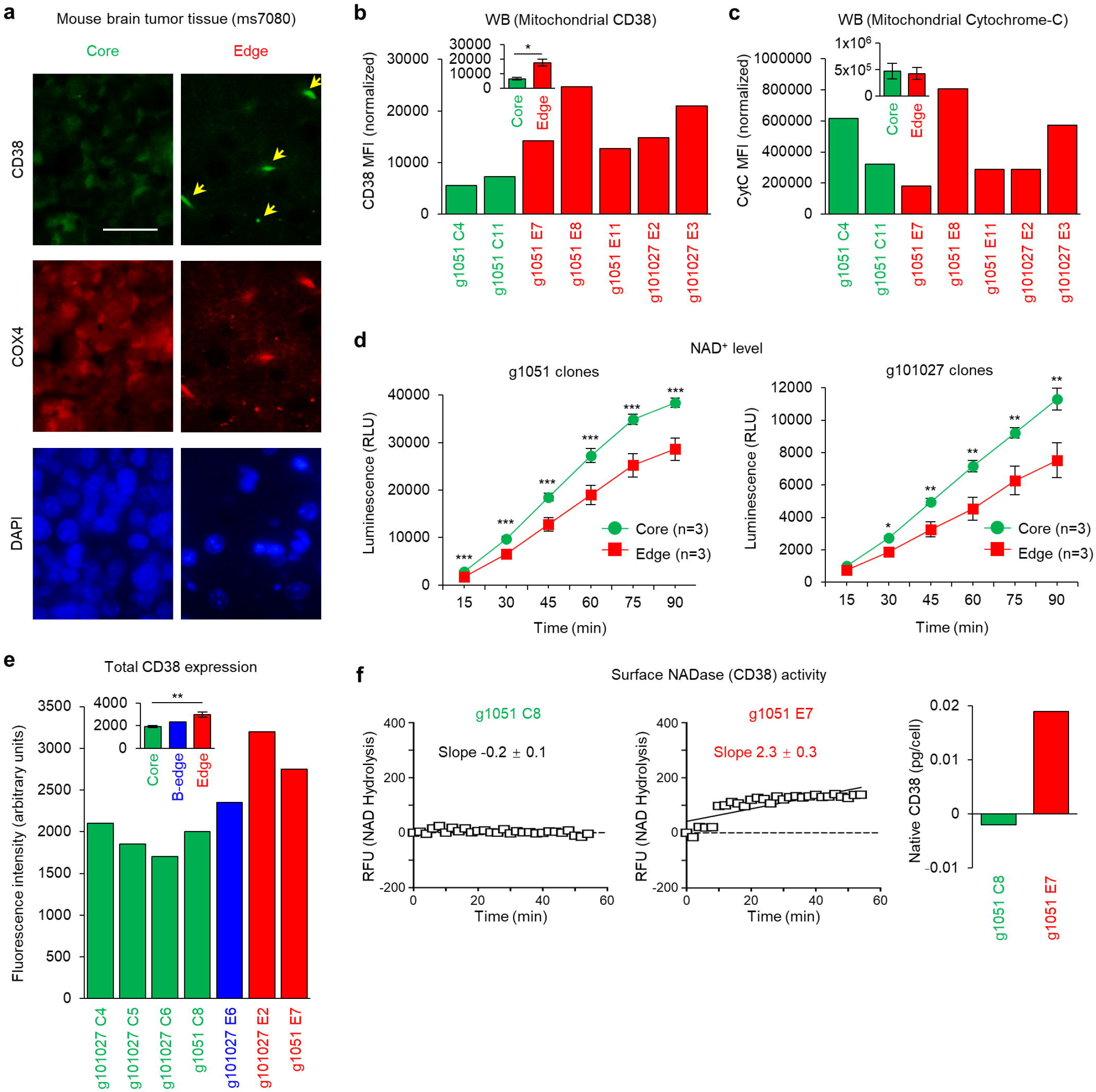
Function of CD38 is mediated within tumor cells’ mitochondria. (**a**) Representative Immunofluorescence (IF) for CD38 (top), COX4 (middle), and DAPI (bottom) of mouse tumors following intracranial injection of mouse glioblastoma cells (ms7080). Scale bar represents 50 μm. Double positive cells for CD38 and COX4 are indicated with yellow arrows. (**b**) Graph indicating fluorescence intensities for mitochondrial CD38 expression of g1051 and g101027 clones. \**p*<0.05. (**c**) Graph indicating fluorescence intensities for mitochondrial Cytochrome-C expression of g1051 and g101027 clones. (**d**) NAD activities in g1051 (top) and g101027 (bottom) Core and Edge clones, measured by luciferase analysis (relative light units, RLU) for qualitative values of NAD^+^ activity. n=3 each. \**p*<0.05, \*\**p*<0.01, \*\*\**p*<0.001. (**e**) Graph indicating fluorescence intensities for total CD38 expression of g1051 and g101027 clones. \*\**p*<0.01. (**f**) Cell surface NADase (CD38) activities in g1051 Core (left; C8) and Edge (right; E7) clones determined by relative fluorescence units (RFUs) versus incubation time in minutes.

### *Intercellular signals between ptEdge cells and astrocytes activates* CD38 *mRNA expression* in culture

Given the high mitochondrial CD38 activities both in tumor cells and astrocytes at the tumor edge, we next examined if ptEdge and astrocytes have any inter-cellular communication for the mutual CD38 activation. ptEdge-derived conditioned media (CM), but not ptCore clones, elevated *CD38* mRNA (up to 3-fold higher) in the normal human astrocyte cell line (NHA) (**Fig. 7a, b**). As a result, we observed that this CM treatment elevates the mitochondria-specific reactive oxygen species (ROS)-related superoxide dismutase (SOD), SOD2, but not the cytosolic SODs (SOD1 or 3) (**Fig. 7c and Extended Data Fig. 7a**). In contrast, the expression levels of reactive astrocyte markers GFAP and LCN2 were unchanged by the same treatment (**Extended Data Fig. 7b**). As reversed trans-cellular signals, we tested if astrocyte-derived trans-cellular signals were capable of influencing *CD38* mRNA expression in ptEdge cells. Pretreatment with NHA-derived CM increased CD38 expression in ptEdge cells; yet the impact was not as prominent (~1.6 fold) as the tumor-to-astrocyte direction (**Fig. 7d, e**). This trans-cellular signal was also observed in the murine tumor model, CD38^high^ tumor cells to mouse WT, but not CD38-null, astrocytes (**Fig. 7f, g**). This data indicates that the activated expression of *CD38* mRNA is likely through the cell-intrinsic mechanism but not by direct inter-cellular transfer of *CD38* mRNA molecule between tumor cells and astrocytes. In these ptEdge CM,4-fold more mitochondrial particles were detected by MitoTracker FACS analysis in comparison to the ptCore counterparts (**Extended Data Fig. 7c-e**), raising a possibility that mitochondrial transfer might be involved in the trans-cellular activation of *CD38* mRNA between Edge cells and astrocytes. Collectively, Edge cells and astrocytes mutually exchange intercellular signals for the activation of *CD38* mRNA expression in the counterparts, at least *in vitro.*

**Fig. 7:**
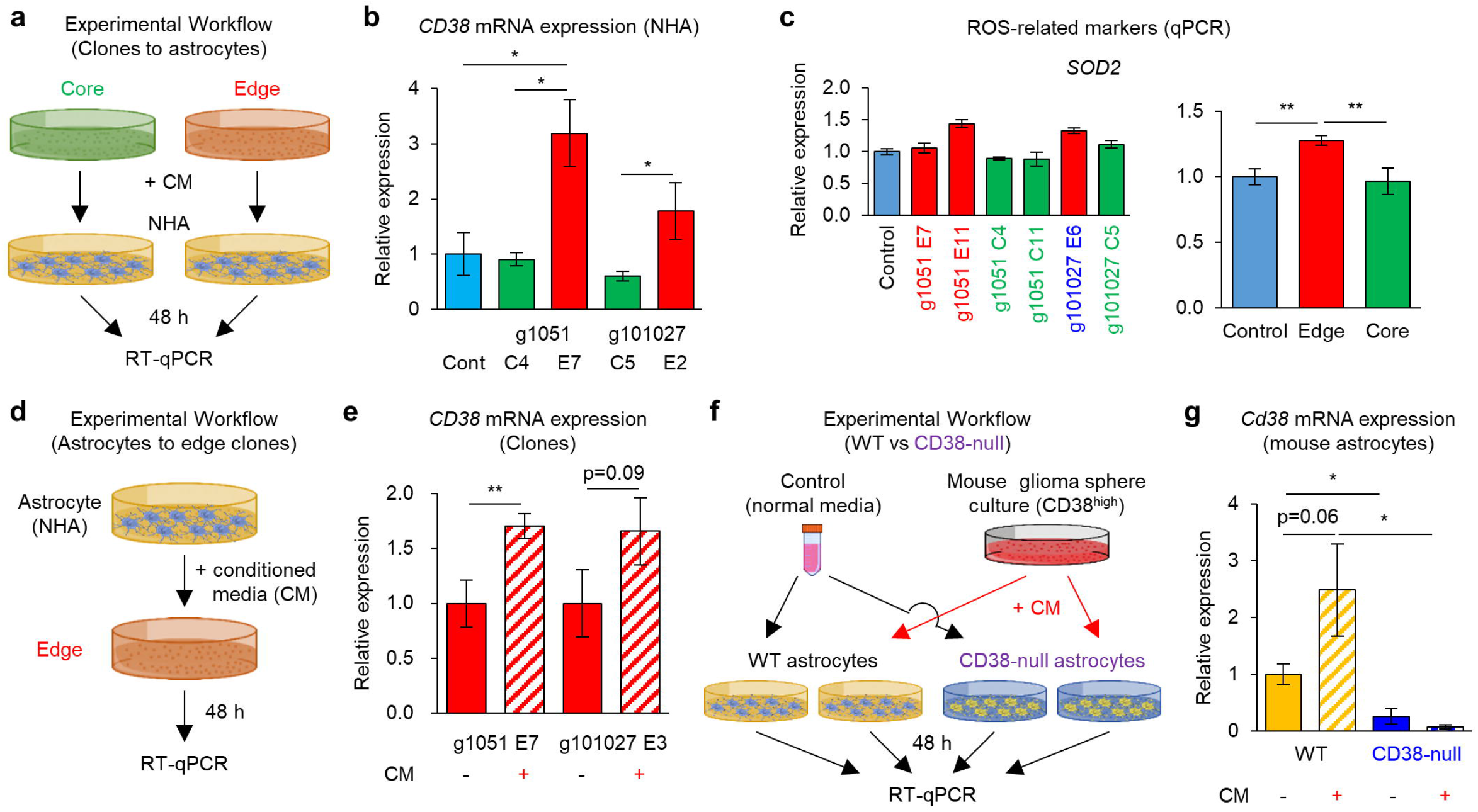
Mitochondrial transfer from ptEdge cells to astrocytes is accompanied with elevation of CD38 *in vitro*. (**a**) Schematic of *in vitro* experiment using the CM from Core and Edge clones to NHAs. (**b**) RT-qPCR analysis comparing relative expression of *CD38* mRNA in CM-treated NHAs. NHAs without CM treatment were used as control (cont). \**p*<0.05. (**c**) RT-qPCR analysis of the expression of ROS-related marker SOD2 in CM-treated NHAs. Data represent the means ± SD (n=3). \*\**p*<0.01. (**d**) Schematic of *in vitro* experimental flow using the CM from NHAs treated onto Edge clones. (**e**) RT-qPCR analysis of the relative expression of *CD38* mRNA in CM-treated g1051 and g101027 Edge clones. \*\**p*<0.01. (**f**) Schematic of *in vitro* experiments using the normal media and CM from WTD glioblastoma cells (CD38^high^) to mouse astrocytes derived from WT and CD38-null mice. (**g**) RT-qPCR analysis of the relative expression of *CD38* mRNA in CM-treated WT and CD38-null mouse astrocytes. \**p*<0.05.

Lastly, towards developing a therapeutic strategy directed at tumor edge lesions, we treated tumor-bearing mice with 78c, a CD38 inhibitor (**Fig. 8a**). We created the tumor-bearing mouse models by injecting tumor core-derived and tumor edge-derived g1051 spheres, followed by systemic treatment with 78c for ten consecutive days. The mice did not exhibit any noticeable adverse events with this treatment. After monitoring the mice for an additional 30 days, the mice were sacrificed to assess edge formation and CD38 expression *in vivo.* We found that the 78c treatment diminished CD38 expression in both edge glioblastoma cells and neighboring astrocytes, accompanied by the substantial inhibition of the edge lesions without noticeable change in the core lesions (**Fig. 8b, c**). These data suggest that pharmacological inhibition of CD38 by systemic administration of 78c can attenuate the formation of edge lesions that retain tumor cell subpopulations to initiate glioblastoma recurrence.

**Fig. 8:**
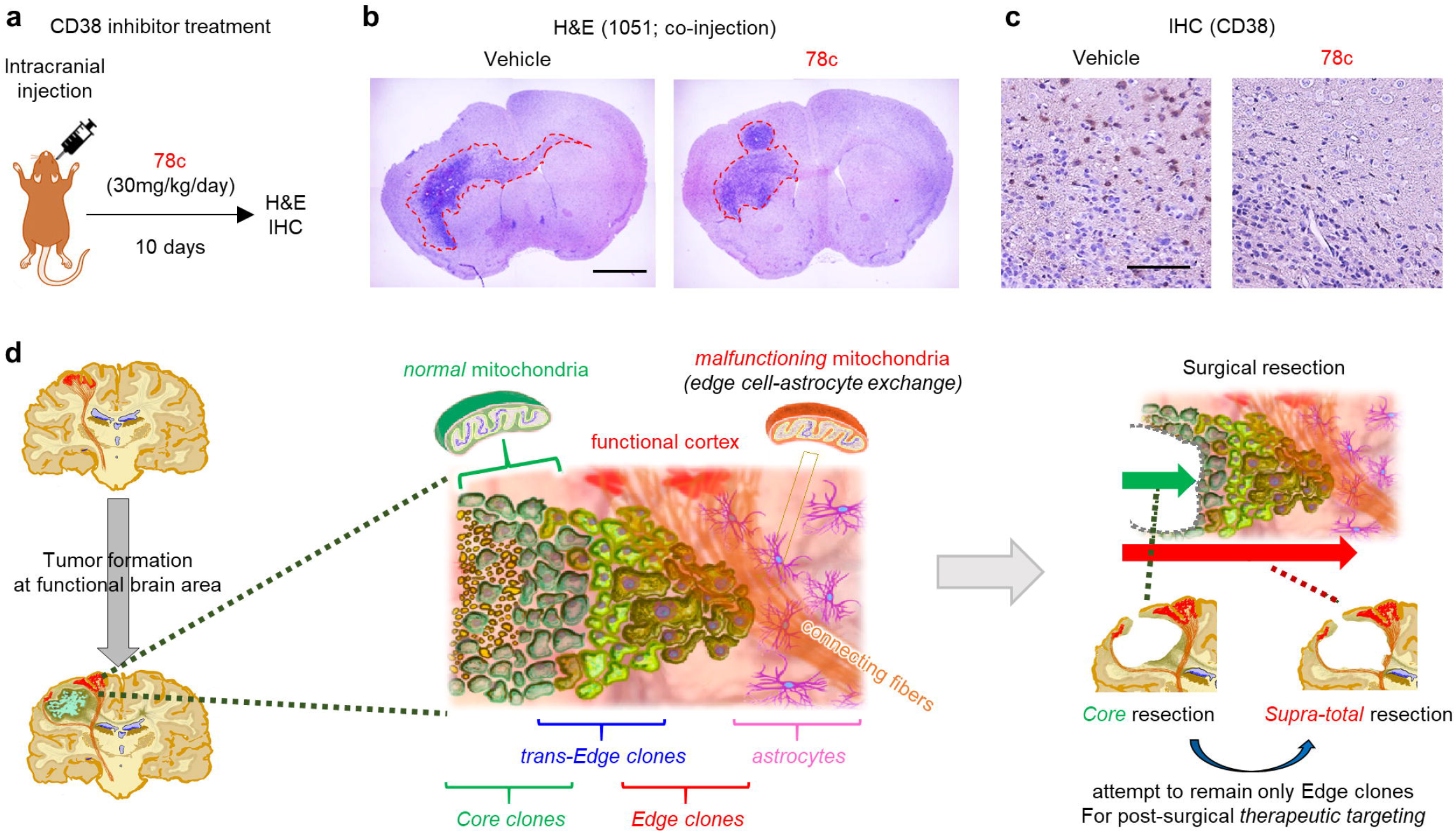
Pharmacological inhibition of CD38 attenuates glioblastoma edge formation. (**a**) Schematic for CD38 inhibitor (78c) treatment on WT mice after tumor challenge. (**b**) Representative H&E staining of Vehicle-treated (left) and 78c-treated (right) WT mouse brains after intracranial injection of ms7080 cells. Scale bars represent 2 mm. (**c**) Representative IHC imaging for CD38 in Vehicle-treated (left) and 78c-treated (right) WT mouse brains. Scale bar represents 100 μm. (**d**) Schematic of Edge to Core progression hypothesis.

## DISCUSSION

The spatial evolution and temporal dynamics of glioblastoma have been studied with various strategies and models, but they remain poorly understood, making effective therapies elusive. In this study, the phylogeny and associated molecular profiling data of clones and tissues indicate a two-step E-to-C progression in glioblastoma (**Fig. 8d**), in part consistent with recent studies analyzing glioblastoma core and edge tissues^3–5^. Unlike the previous studies, the present data elucidate the isolated tumor cell-specific pathobiology, yet did not determine whether the tumor edge clones, particularly ptEdge, accumulate at the SVZ and/or corpus callosum. Given the infiltrating pattern of glioblastoma in mouse brains resembled patient tumors with preferential accumulation at the SVZ and corpus callosum, both lesions are possibly the dominant sources of these edge-derived clones, although validation is required. From a clinical standpoint, it remains to be proven whether ptEdge clones are capable of evolving into trans-ptEdge and subsequently to ptCore clones *in vivo,* especially in response to radiation and chemotherapy. Treatment-naive tumor edge and core cells express distinct key transcription factors^4^, activated kinases^6^ and metabolites^9^. Post-surgical selection pressure by chemo-radiation causes secondary dynamic changes to these molecules in the remaining tumor edge cells, particularly in those that initiate tumor (core) recurrence/relapse. These molecular drivers may determine the fate of the distinct tumor clones, as well as trigger their conversion^19^ and intercellular communication^7^ to promote both glioblastoma heterogeneity and therapy resistance. Among the three clonal subpopulations identified, the trans-ptEdge cells retained the *‘edgeness’* phenotype, while also becoming core-like, albeit through unknown molecular mechanisms. According to a recent paper by Lin CM *et al.,* transient and reversible peripheral hypoxia in a mouse cell line model was associated with astrocytic activation and therapy resistance due to vascular malfunction^23^. Based on the data with the slice culture models, trans-ptEdge clones were only loosely connected with brain vasculature, in sharp contrast to ptEdge clones, which had strong affinity for vasculature. If this assay faithfully reflects the phenotype of tumor cells in the brains of patients, our results suggest that transient clones might be primed at the initial phase to adapt to the more hypoxic and acidic challenges of the adjacent core microenvironment as the tumor expands. Notably, these tumor cells were derived from the edge of the tumors. Secondary tumor challenge, at least in mouse brains without any therapy-induced pressure, resulted in the formation of new core lesions in addition to the edge. In principle, prevention of the conversion of the ptEdge to trans-ptEdge clones would be one therapeutic venue that should inhibit the development of recurrent tumor core. Future studies are required to uncover how these transitional clones undergo their spatiotemporal phenotypic shift by acquiring adaptive molecular profiles.

Among the identified clones, ptEdge exhibited substantially lower NAD^+^ levels accompanied by elevated activity of the NADase CD38. Both astrocytic and tumor CD38 were mandated for the formation of tumor edge lesions, directly affecting overall tumor aggressiveness. Tumors with disrupted CD38 production might have lost the edge-forming tumor cells, although it is also plausible that CD38-removed edge-forming cells, at least a subset of them, might have *differentiated* into core cells through an E-to-C-related mechanism. NAD^+^ can function as a double-edged sword, providing energy for somatic cells as well as cancer cells including glioblastoma^24,25^. Elevated NAD^+^ activity in edge clones may subsequently promote their relocalization to the core, elevating overall tumor aggressiveness. This hypothesis warrants experimental scrutiny in the future. In addition, exchange of trans-cellular signals through mitochondrial particle-rich CM induced *CD38* mRNA expression both in edge cells and astrocytes in a cell-intrinsic manner. Yet, the mechanism of action for *CD38* activation remains undetermined. Further experimentation is required to understand whether this phenotype also occurs *in vivo* and how it is molecularly regulated.

For therapeutic development, CD38 targeting *via* systemic administration of 78c presented promising results, but this treatment may also induce (partial) core differentiation in exchange for reducing edge lesions. Thus, subsequent combinatorial therapies with core-targeting interventions might be taken into consideration. Future studies are also warranted to determine whether the fine tuning of NAD^+^ activity within the tumor edge ecosystem and at subsequent core lesions would make any clinical impact in glioblastoma and other cancers.

## Acknowledgements

We would like to express our sincere appreciation to all the patients and families, who kindly allowed us to obtain their tumor samples for this study. To achieve the best benefit for the neuro-oncology research on behalf of our patients, we would be more than willing to share our patient-derived experimental models, described in this study with any scientists who express interest, once the appropriate paperwork is agreed and completed in place. We would also thank all our collaborating scientists, as well as the assigned reviewers and Editor for this manuscript, for constructive comments and suggestions. We acknowledge the contribution by all the members in the Nakano and Lund laboratories (past and present) for technical help. We also deeply thank Oriental Yeast Co., Ltd. for supply of NMN for this project. This work was supported by NIH grants R01NS083767, R01NS087913, R01CA183991, and R01CA201402 to IN. MG was supported by the NIA IRP, NIH.

## Author Contributions

Leading conceptualization of the study: IN. Financial support: IN, FL. Overall design of the study: DY, IN, with input from TK, HIK, DN, ZG, FEL. Laboratory practice: DY, DB, HJC, XG, SO, VLF, IS, SY, FZ. Analysis of data; DY, DB, HJC, XG, MG, SY, HZ, DKC, NZ, JS. Establishment of NSCL61 cells: TK. Drafting the article: DY, MAN, IN. Critical revision of the article: MG, IN. All authors had substantial input to the logistics of the work and revised and approved the final manuscript. The authors know their accountability for all aspects of the study ensuring that questions regarding the accuracy and integrity of any part are appropriately investigated and resolved. The corresponding author had full access to all of the data and the final responsibility to submit the publication.

## MATERIALS AND METHODS

### Ethics

All work related to human tissues was performed under an institutional review board (IRB)-approved protocol compliant to the National Institute of Health (NIH) guidelines (IRB approval number: N151013001). For the pre-clinical studies, 32 patient-derived glioma sphere models were used, including two pairs of tumor core- and edge-derived clones (1051E and C, 101027E and C) as well as one tumor core-derived sphere line (301016C). The senior author (IN) performed supra-total resection of glioblastoma tumors under the awake setting and resected both tumor core (T1-Gadolinium(+) tumors) and edge (T1-Gadolinium(-)/T2-FLAIR abnormal tumors in the non-eloquent deep white matter) to achieve maximal tumor cell eradication without causing any permanent major deficit in the patients (see **Extended Data Fig. 1a, b** for the definition of tumor core and edge). After confirmation that sufficient tumor tissue from both lesions were secured for clinical diagnosis, residual tissues were provided to the corresponding scientists following de-identification of patient information. Both the core-derived and edge-derived glioma spheres were established in the same culture condition, and their spatial identities *(‘coreness’* and *‘edgeness’)* were confirmed by xenografting experiments in mouse brains (details in Ref. 7). Only those that passed this confirmation were used for this study. The other patient-derived glioma sphere models were established as “core-associated glioma spheres” using the same protocol and reported elsewhere^5^. All the patient-derived glioma models were periodically checked for mycoplasma contamination and Short-Tandem-Repeat (STR) profiling (DNA fingerprinting) for identification of the patient samples.

### Data reliability and validity

All the experiments were performed by two or more researchers involved in each procedure. The experiments using cell culture were repeated at least three independent times to obtain consistent data. All the raw data are placed in **Extended Data Fig. 8**. The corresponding author (IN) confirmed the agreement of everything related to this study with all the involved authors.

### Statistical analyses

Statistical analysis was performed using XLSTAT 2018.5, SPSS statistical package version 25, and Graphpad Prism 7.0 software. All data were presented as the mean ± SD. P-values <0.05 were considered statistically significant. Statistically significant differences in Kaplan-Meier survival curves were determined by log-rank analysis.

### Data availability

Data are available in the Gene Expression Omnibus (GEO) under accession GSE135408.

### Code availability

All custom code used in this work is available from the corresponding authors upon reasonable request.

## SUPPLEMENTARY MATERIALS

### Murine glioblastoma models

Glioma sphere cultures were established from murine tumors formed by *Cre; Nf1*^f/+^; *p53*^f/f^; *Pten*^f/+^ mice (ms7080), and by transforming p53-deficient neural stem cells (NSC) with oncogenic HRas^L61^ (NSCL61).

### Cell culture

Both patient-derived and murine glioma spheres were cultured in the defined medium containing DMEM/F12/ Glutamax (Invitrogen) supplemented with B27 (Miltenyi Biotec), heparin (2.5 mg/ml), basic fibroblast growth factor (bFGF) (Peprotech, 20 ng/ml), and epidermal growth factor (EGF) (Peprotech, 20 ng/ml). Growth factors (bFGF and EGF) were added twice a week and the culture medium was changed every 7 days. Details are described in our previous studies^3,5–9,26^.

### Brain slice culture

Mice at the age of postnatal day 7 were rapidly sacrificed and the heads were briefly placed in 70% ethanol for dissection of the brains. Under aseptic conditions, 350 μm-thick whole-brain (sagittal or coronal) sections were cut and collected in sterile medium. Patient-derived core and edge tumor cells (labeled with GFP- or mCherry-carrying lentivirus) were seeded on the brain slices (10^5^ cells) along with 5 μl matrix gel. Brain slices were maintained at 37°C and 5% CO_2_ for 2 days. Subsequently, the brain slices were fixed for 8 h at 4°C in 4% PFA and stained with collagen4 (Millipore, AB756P) to label the brain vasculature with anti-rabbit Alexa555 as second antibody. Images and videos were captured with Nikon A1 confocal microscope.

### RNA isolation and Reverse Transcription followed by Quantitative PCR (RT-qPCR) analysis

Total RNA was extracted using the RNeasy mini kit (QIAGEN) according to the manufacturer’s instructions. RNA concentration was determined using Nanodrop One (Thermo Scientific). cDNAs were synthesized using iScript reverse transcription supermix (Bio-Rad) according to the manufacturer’s protocol. RT-qPCR analysis was performed on StepOnePlus thermal cycler (Thermo scientific) with SYBR Select Master Mix (Thermo scientific). *GAPDH/Gapdh* mRNA was used as an internal control. Details are described in our previous studies^3,5,7,8,19^ Primer sequences for the genes tested in this study are shown in Supplementary Table 1.

### Immunohistochemistry (IHC)

IHC was performed as previously described and signals were detected using the DAB substrate kit (Vector). For double staining, donkey IgG H&L (alkaline phosphatase) pre-adsorbed antibody (Abcam) was used to detect primary antibodies and detected by the liquid fast-red substrate kit (abcam). Primary antibodies used in this study recognized CD38 (Santacruz), GFP (Abcam), mCherry (Abcam), or CoxIV (Cell Signaling). Details are described in our previous studies^3,5,7,8,19^

### Immunofluorescence (IF)

IF was performed as previously described and cells were probed with following primary antibodies: anti-CD38 (Santa Ccruz) and anti-COX IV (Cell Signaling). Alexa 488- and 594-labeled secondary antibodies were used to visualize immunofluorescence signals. Representative images were photographed using confocal microscopy (Zeiss)^19^.

### Flow cytometry

Surface CD38 expression on human and murine cells was analyzed using a FACSCanto II (BD Biosciences) flow cytometer. Cells were filtered through a 70-μm nylon cell strainer (BD Biosciences), washed and incubated with staining media (PBS containing 2% donor calf serum and 2 mM EDTA) for 20 min on ice with either an anti-mouse antibody staining mix consisting of LIVE/DEAD™ Aqua dead cell stain (dilution 1:800 – ThermoFisher Scientific), 10 μg/ml FcBlock (clone 2.4G2 – BioXCell) and CD38-PE-Cy7 (clone 90, dilution 1:200 – eBioscience), or an antihuman antibody staining mix consisting of LIVE/DEAD™ Aqua dead cell stain (dilution 1:800 – ThermoFisher Scientific), 2.38 μg/ml Human Gamma Globulin (Jackson ImmunoResearch) and CD38-PE-Cy7 (clone HIT2, dilution 1:1200 – eBioscience). Following each staining, the cells were washed and resuspended with staining media prior to flow cytometric analysis. For mitochondrial particle analysis, conditioned media were collected from multiple clones labeled with MitoTracker Red CMXRos and were used to sort labeled mitochondrial fraction by cell sorter configured with 561-nm air-cooled laser. Analysis of the flow cytometry data was performed according to the methods described in our previous studies^3^.

### Surface CD38 activity assay

The NAD glycohydrolytic activity of surface CD38 on murine and human cells was measured using a sensitive, continuous, fluorometric assay platform that used 1,N^6^-ethenonicotinamide adenine dinucleotide (εNAD; MilliporeSigma) as a substrate. Briefly, 1 x 10^6^ cells were incubated in duplicate wells with 25 μM εNAD in a serum-free 150 μl reaction volume. The appearance of the fluorescent εADP-ribose reaction product was measured every 2 minutes for a total of 45-60 min at excitation (λ_exc_) and emission (λ_em_) wavelengths of 310 nm and 410 nm, respectively, and at 37 °C using a SpectraMax M2 microplate reader (Molecular Devices). CD38 enzymatic activity was plotted as relative fluorescence units (RFUs) versus incubation time in minutes, and the slope values were calculated by linear regression.

### Western blot analysis

CD38 protein levels in whole-cell lysates were quantified by western blotting, according to the protocol described in our previous studies^3,5,7,19^ Briefly, 10 μg of total protein was subjected to SDS-PAGE on 4-15% (w/v) gradient gels, and the protein was electroblotted onto nitrocellulose membranes. The membranes were first stained for total protein using the REVERT™ Total Protein Stain solution (Li-COR). The membranes were then blocked for 1 h at room temperature in Odyssey^®^ blocking buffer (OBB, Li-COR) diluted 1:1 with TBS, and incubated overnight at 4 °C with anti-CD38 rabbit monoclonal antibody (Abcam) at a 1:3,000 dilution. Following the primary antibody incubation, the membranes were washed 3X (10 min each) with TBS containing 0.1% Tween-20 (TBS-T), and incubated for 1 h at room temperature with anti-rabbit IgG (H+L) secondary antibody conjugated to IRDye 800CW (Li-COR) at a dilution of 1:30,000. The membranes were washed 3X (10 min each) with TBS-T, and scanned and analyzed using an Odyssey^®^ IR scanner (Li-COR). Total protein and CD38 protein levels were detected in the 700 nm and 800 nm channels, respectively.

### Measurement of NAD^+^/NADH

The levels of NAD^+^ and NADH were measured by the NAD/NADH-GloTM Assay (Promega) following the manufacturer’s protocol. Briefly, 1×10^5^ cells were lysed with 400 μl of PBS and then incubated with 400 μl of 0.2 N NaOH with 1% Dodecyltrimethylammonium bromide (Sigma-Aldrich). To measure NAD^+^, equal volumes of 0.4 N HCl were added to the lysed cell sample (400 μl), followed by heating at 60°C for 15 min. After incubation at room temperature for 10 min, each sample was incubated with 0.5 M Trizma base buffer (200 μl, Sigma-Aldrich). To measure NADH, the lysed cell samples (400 μl) were heated at 60°C for 15 min, left at room temperature for 10 min, followed by incubation with equal volume of HCl/Trizma solution. Finally, samples were seeded into 96-well plates for NAD/NADH-Glo detection reagent for 30 min. Data were compared with the DMSO-treated control cells.

### Quantification of mtDNA copy numbers

The relative levels of mtDNA copy number were measured by qPCR-based assay using human mitochondrial DNA copy number assay kit (The Detroit R&D, Inc.) following the manufacturer’s instructions. All quantitative real-time PCR reactions were performed using a LightCycler 480 Instrument (Roche Life Science). In brief, purified genomic DNA (15 ng) was amplified with primers targeting mitochondrial DNA and nuclear DNA in separate 20-μL PCR reactions containing 10 μL of rtPCR reaction mix, and 500 nmol/L of forward and reverse primers. The thermal profile included an initial denaturation step of 10 min at 95°C, followed by 40 cycles of 95°C for 15 s and 60°C for 60 s. For each sample, mtDNA and nDNA targets were run in duplicate. Results from duplicates were averaged to compute mean Ct values for both targets. Their difference (ΔCT) were then normalized to that of a positive control sample measured on the same plate by using ΔΔCT method to obtain relative mtDNA copy number.

### Oxygen Consumption Rate (OCR)

OCR in each sample was measure according to the protocol described in our recent studies^7,26^. Briefly, XFE-24 Seahorse plates (Seahorse Biosciences, 102340) were coated with 50 μL laminin (Sigma) diluted in PBS (10 μg/mL) overnight. The next day, each wells was washed three times with PBS, and cells were plated at a density of 1.0×10^5^ cells per well. These cells were cultured overnight in phenol-free DMEM media to adhere on culture dishes for 24 hours at 37°C and 5% CO_2_. The Seahorse 24 optical fluorescent analyzer cartridge was prepared in the interim by adding 1 μM oligomycin, 1 μM FCCP, and 0.5 μM rotenone to each cartridge port. OCRs (pmole/min) and extracellular acidification rate (mpH/min) were then measured for each sample at 37°C using the Seahorse Bioanalyzer instrument, according to the company’s protocol.

### Animal experiments

All animal experiments were performed under an Institutional Animal Care and Use Committee (IACUC)-approved protocol according to NIH guidelines (#20295). C57BL/6 mice and immunocompromised mice (SCID Beige) were purchased from Charles River and CD38 KO mice were purchased from the Jackson Laboratory. For intracranial injection, mice were anesthetized with ketamine/xylazine and fixed in place and dissociated BC cells were stereotactically injected into the striatum of mice. Detailed protocols are fully described in our studies^3,5,7,8^.

### CD38 inhibitor (78c) treatment

Systemic administration with 78c to C57BL/6 mice was performed by intraperitoneal injection (i.p., 15 mg/kg/dose) twice daily for 10 consecutive days. Control mice received vehicle (5% DMSO, 15% PEG400, 80% of 15% hydroxypropyl-γ-cyclodext⋂n (in citrate buffer pH 6.0)) injections. Treated mice were closely monitored for their systemic condition following treatment.

### RNA sequencing

Detailed protocol for RNA sequencing is described in our studies^3,5,7,9^ Briefly, isolated RNA samples were sequenced commercially at QUICK BIOLOGY (http://www.quickbiology.com). Following depletion of ribosomal (r)RNA, libraries were prepared and sequenced using an Illumina HiSeq 4000 instrument, PE150, for a total of 80 million reads per sample.

### RNA-Seq analysis

As described in our previous studies^3,5,7,9^ STAR (version 2.5.3a) was used to align the raw RNA-Seq fastq reads to the mouse reference genome (GRCm38 p4, Release M11) from Gencode with parameters --outReadsUnmapped Fastx; --outSAMtype BAM SortedByCoordinate; --outSAMAttributes All^27^. Following alignment, HTSeq-count (version 0.9.1) was used to count the number of reads mapping to each gene with parameters -m union; -r pos; -t exon; -i gene_id; -a 10; -s no; -f bam^28^. Normalization and differential expression were then applied to the count files using DESeq2.

### Systems biology analysis

For generating networks, a dataset containing gene identifiers and corresponding expression values were uploaded for Ingenuity Pathway Analysis. Each identifier was mapped to its corresponding object in Ingenuity’s Knowledge Base. A fold change cutoff of ±2 and p-value < 0.05 was set to identify molecules whose expression was significantly differentially regulated. These molecules, called network-eligible molecules, were overlaid onto a global molecular network developed from information contained in Ingenuity’s Knowledge Base. Networks of network-eligible molecules were then algorithmically generated based on their connectivity. The functional analysis was used to identify the biological functions and/or diseases that were most significant to the entire data set. Molecules from the dataset that met the fold change cutoff of ±2 and *p*-value < 0.05 and were associated with biological functions and/or diseases in Ingenuity’s Knowledge Base were considered for the analysis. Right-tailed Fisher’s exact test was used to calculate a *p*-value determining the probability that each biological function and/or disease assigned to that data set is due to chance alone. For the Gene Set Enrichment Analysis (GSEA) and Gene Ontology enrichment analysis (GO), the following websites were used: http://software.broadinstitute.org/gsea/index.jsp and http://geneontology.org/, respectively.

### DNA sequencing

The Agilent SureSelect Human All Exon V6 Kit was used to capture DNA fragments for whole-exome sequencing. Following this sequencing process, paired-end 150 bp reads were generated with an Illumina HiSeq2500. The sequenced reads from DNA sequencing were mapped to hg19 using the Burrows-Wheeler Aligner (BWA)^29,30^.

### DNA mutation calling and copy number alteration analysis

MuTect and SomaticIndelDetector (GATK) were used to detect single-nucleotide variants and indels, and the called mutations were annotated using Variant Effect Predictor (VEP)^31–33^. For the clone samples without matched blood samples (g1051 clones), possible somatic mutations were filtered using an in-house normal variant pool, consisting of variants detected in our inhouse blood pool. For phylogenetic tree generation of g101027 clones, Phylip was used using all called somatic mutations with variant allele frequency (VAF) >= 0.05 and altered reads >=5. For g1051 clones, only exonic events with VAF >= 0.05 and altered reads >=5 were obtained from previous steps, and these events were subjected to Phylip to generate phylogenetic tree. ngCGH was used to estimate copy number alterations of all g101027 and g1051 clones, and g101027 blood sample was used as control (normal) sample.

### Computational assessment of cancer-associated mutations

Clones derived from the edge and core tissues are sequenced and analyzed individually. For each set of full exome sequence data, ENTPRISE^34^ and ENTPRISE-X^35^ were applied to identify mutations associated strongly with pathogenicity. From these sample-associated mutations, cancer-driver targets by cross-checking the COSMIC database^36^ were selectively investigated and specifically identified. Only mutations found in the cancer-driver targets as indicated by COSMIC were retained for further investigation.

### Mass Spectroscopy for Metabolomics

Approximately 1 mg of human glioblastoma tissue or 10^6^ cells were collected for analysis. Resected tissue was classified as originated from the tumor core or edge. The sample was pulverized with pestle and immediately placed in ice cold methanol. After homogenization, 4 ml ice cold methanol was added and the sample was centrifuged at 10,000 rpm at 4°C for 10 min. Extracted metabolites were analyzed by flow injection-time of flight mass spectrometry on an Agilent 6550 QTOF instrument operated in the negative mode as previously described^37^. Detectable ions were putatively annotated by matching measured mass-to-charge ratios with theoretical masses of compounds listed in the human metabolome database v3.0 using a tolerance of 0.001 ūamu^38^. Pathway definition of differentially abundant metabolites was performed with the Small Molecule Pathway Database^39^. For comparison, *p*-values were calculated by two-tailed, heteroscedastic *t*-test and were adjusted for FDR according to the Benjamini-Hochberg procedure. All calculations were done in Matlab.

## Supplementary Figure Legend

**Extended Data Figure 1.**

(**a**) Representative MRI of another glioblastoma patient (case 2) depicting the glioblastoma core area (red) and edge areas (yellow). V: lateral ventricle; SC: subcortical edge; SVZ: subventricular zone.

(**b**) Representative MRI and surgical view of another glioblastoma patient (case 3).

(**c**) Representative images of fluorescence of labeled ptCore and ptEdge spheres.

(**d**) Representative H&E staining of mouse brains harboring g101027 ptCore and ptEdge tumor cells 2 days after intracranial injection. Scale bars represent 2 mm (left) and 200 μm (right).

(**e**) Representative IHC imaging for GFP (top) and mCherry (bottom) in msCore (left) and msEdge (right) 2 days after intracranial injection of g101027 tumor cells. Scale bar represents 100 μm.

**Extended Data Figure 2.**

(**a**) Experimental design for establishment of g1051 Edge and Core clones.

(**b**) Gene expression panel comparing Core and Edge clones’ single nucleotide polymorphisms (SNP) in key upregulated genes in glioblastoma.

(**c**) Representative IHC for human mitochondria to label tumors derived from g101027 Core (left), trans-Edge (middle), and Edge (right) clones. Injection site indicated in yellow asterisks. Red arros indicate the infiltrating tumor cells within corpus callosum. Scale bar represents 1 mm.

(**d**) Hierarchal clustering for g1051 (left) and g101027 (right) clones.

(**e**) Graphical representation of DNA mutation positions in g1051 (left) and g101027 (right) clones (n=9 and 6, respectively).

(**f**) Panels of major gene mutations in g1051 (left) and g101027 (right) clones.

(**g**) Gene mutation panel comparing common gene mutations in distinct g1051 (left) and g101027 (right) clones.

**Extended Data Figure 3-1.**

(**a**) Heatmap of RNA-seq data comparing ptCore (n=3), ptEdge (n=3) and normal tissue (n=3) gene expressions.

(**b**) Heatmap of RNA-seq data from g1051 Core, trans-Edge, and Edge clones (n=15).

(**c**) Heatmap of RNA-seq data from g101027 Core, trans-Edge, and Edge clones (n=6).

(**d**) IPA demonstrating elevation of Myc and HIF1A pathways in ptCore tissues.

(**e**) IPA demonstrating elevation of Myc and HIF1A pathways in ptCore clones from g1051 and g101027 (n=15 in total).

(**f**) GO analysis of RNA-seq data for comparison between ptCore and ptEdge tissues.

**Extended Data Figure 3-2.**

(**g**) Other GSEA plot of the elevated pathways including those related to inflammation, glycolysis, and epithelial-mesenchymal transition (EMT) in ptCore tissues.

(**h**) MetaCore analysis demonstrating elevation of c-Myc pathway in ptCore tissues.

(**i**) MetaCore analysis demonstrating elevation of c-Myc and HIF1 pathways in Core clones.

(**j**) Unbiased heatmap comparing metabolite expressions in g1005, g1014 and g1057 ptCore and ptEdge tissues (n=36 in total).

(**k**) Volcano plot of metabolite data comparing ptCore and ptEdge tissues.

**Extended Data Figure 3-3.**

(**l**) Heatmap plot of metabolite data comparing g1051 (top) and g101027 (bottom) clones (n=9 and 9, respectively).

(**m**) Volcano plot of metabolite data comparing g1051 (top) and g101027 (bottom) clones.

(**n**) Pathway enrichment analysis of metabolites in g1005, g1014 and g1057 ptCore and ptEdge tissues.

**Extended Data Figure 3-4.**

(**o**) Metabolic map comparing relative expressions of metabolites in g1057 Edge and Core tissues.

**Extended Data Figure 3-5.**

(**p**) Metabolic map comparing relative expression of metabolites in g1051 (top) and g101027 (bottom) clones.

**Extended Data Figure 3-6.**

(**q**) Mitochondrial DNA (mtDNA) copy number by mtDNA seq using g101027 Core and Edge clones (n=12).

(**r**) Mitochondrial respiration in g101027 Core and Edge clones, measured by oxygen consumption rate (OCR).

(**s**) Proton leak and ATP production in indicated g1051 clones.

**Extended Data Figure 4.**

(**a**) Volcano plot of RNA-seq data comparing ptCore and ptEdge tissues, highlighting CD38 (arrow) in Edge tissues.

(**b**) Volcano plot of RNA-seq data comparing normal brains and ptEdge tissues, highlighting CD38 (arrow) in Edge tissues.

(**c**) Bar graph comparing relative expression of CD38 by RNA-seq of ptCore, ptEdge, and normal brain tissues. \*\*\**p*<0.001.

(**d**) Single-cell (sc) RNA-seq analysis comparing *CD38* mRNA expression in distinct cell types in mouse normal brain.

(**e**) FACS for cell surface CD38 expression in wildtype (WT) and CD38-null astrocytes.

(**f**) Cell surface NADase activities in WT and CD38-null astrocytes, measured by relative fluorescence units (RFUs) versus incubation time in minutes.

**Extended Data Figure 4-2.**

(**g**) GSEA plots of the additional upregulated pathways in WTD and KOD tumors.

(**h**) GSEA plot of the additional upregulated pathways in WT and CD38 KO cortical tissues.

**Extended Data Figure 6.**

(**a**) Western blotting for mitochondrial CD38 expression of g1051 and g101027 clones.

(**b**) Western blotting for nuclear CD38 expression of g1051 and g101027 clones.

(**c**) Graph indicating fluorescence intensities for nuclear CD38 expression of g1051 and g101027 clones.

(**d**) Representative cell surface CD38 expression in g1051 Core (top; C8) and Edge (bottom; E7) clones, determined by FACS.

(**e**) FACS analysis of cell-surface CD38 expression in WTD (left) and KOD (right) glioblastoma cells.

(**f**) NAD activity in individual g1051 clones measured by luciferase analysis (RLU).

(**g**) Total CD38 expression of g1051 and g101027 clones, determined by Western blotting.

(**h**) Cell surface NADase (CD38) activities in g101027 clones determined by relative fluorescence units (RFUs) versus incubation time in minutes.

**Extended Data Figure 7.**

(**a**) RT-qPCR analysis of the expression of other ROS-related markers (SOD1 and SOD3) in CM-treated NHAs. Data represent the means ± SD (n=3). \**p*<0.05, \*\**p*<0.01.

(**b**) RT-qPCR analysis of the relative expression of reactive astrocyte markers in CM-treated NHAs. Data represent the means ± SD (n=3). \**p*<0.05, \*\**p*<0.01, \*\*\**p*<0.001.

(**c**) Schematic of the collection of CM from g1051 Core (C4) and Edge (E12) clones.

(**d**) Representative FACS images for MitoTracker Red using the conditioned media (CM) from Core (left) and Edge (right) clones.

(**e**) Graph indicating the number of mitochondrial particles in g1051 Core (C4) and Edge (E12) clones.

**Extended Data Figure 8.**

(**a**) Original western blotting gel for mitochondrial CD38 expression of g1051 and g101027 clones.

(**b**) Original western blotting gel for mitochondrial Cytochrome-C expression of g1051 and g101027 clones.

(**c**) Original western blotting gel for nuclear CD38 expression of g1051 and g101027 clones.

