## Supplemental Figures for "Spatial heterogeneity of glioblastoma cells reveals sensitivity to NAD^+^ depletion at tumor edge"

#### Extended Data Figure. 1

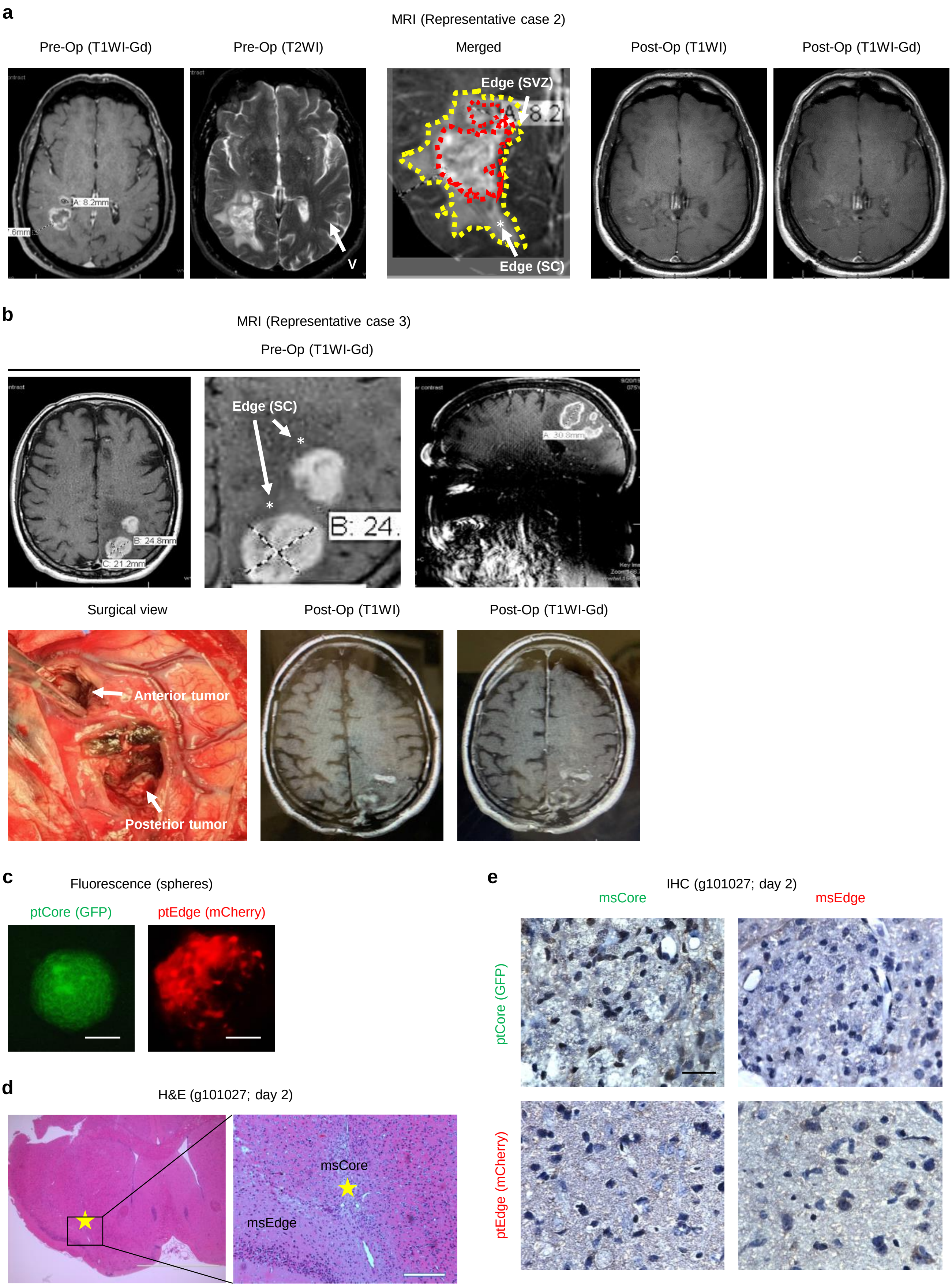

### Extended Data Figure. 2-1

**a**

Establishment of clones (g1051 clones)

| Seeded number/well | Wells w/ spheres (Core) | Wells w/ spheres (Edge) |
|---|---|---|
| 30 | 0/66 | 0/66 |
| 50 | 0/66 | 0/66 |
| 100 | ~20/132 | ~20/132 |

Core clones: 11
Edge clones: 13

~30 Passages for 10 months

**b**

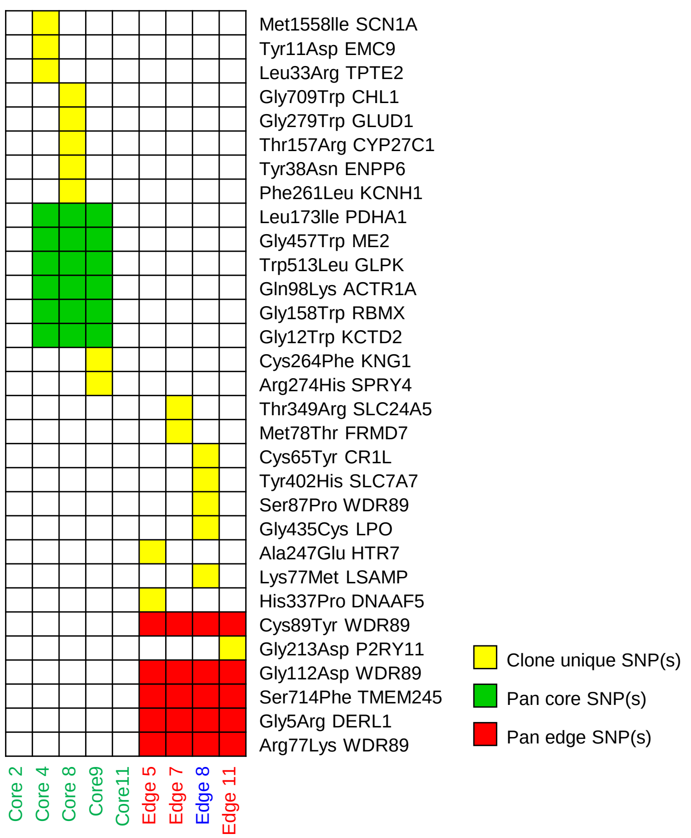


**c**

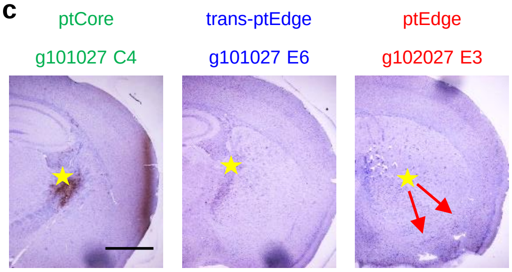


**d**

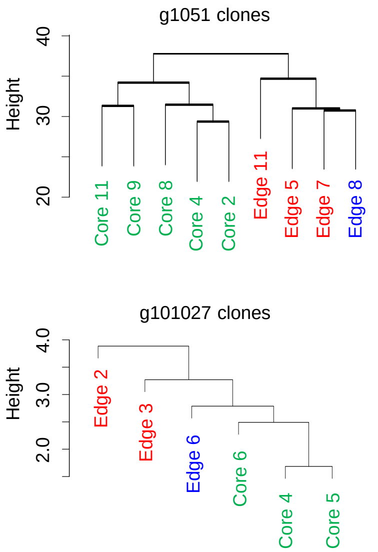


**e**

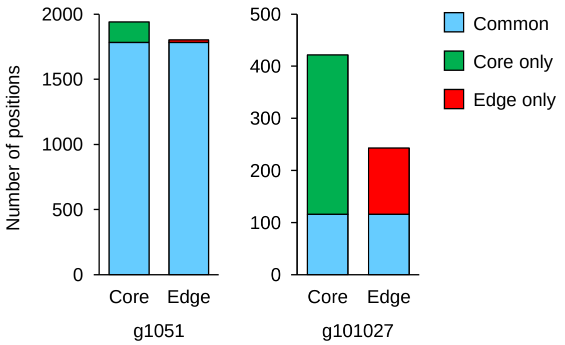

#### Extended Data Figure. 2-2

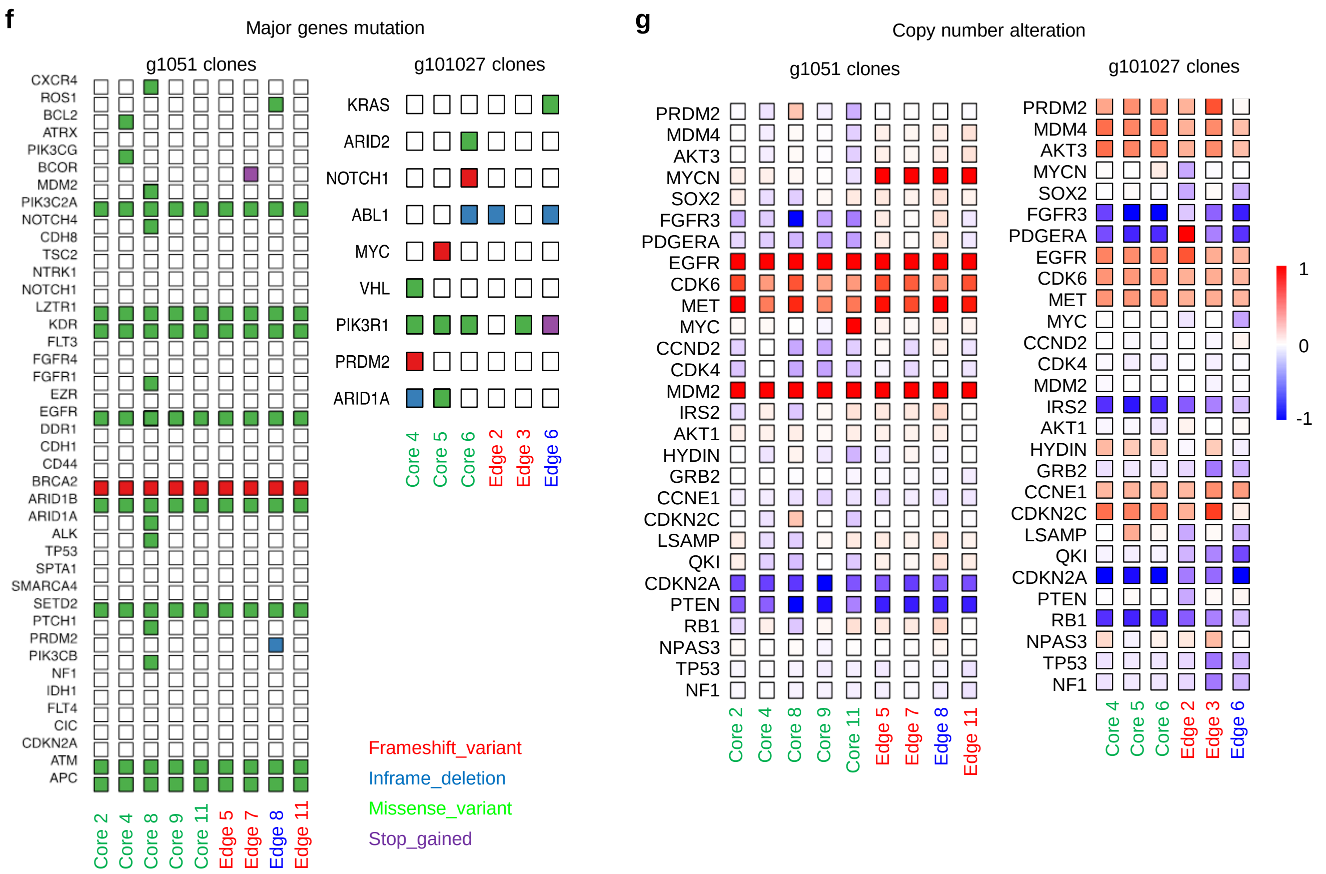

**Extended Data Figure. 3-1**

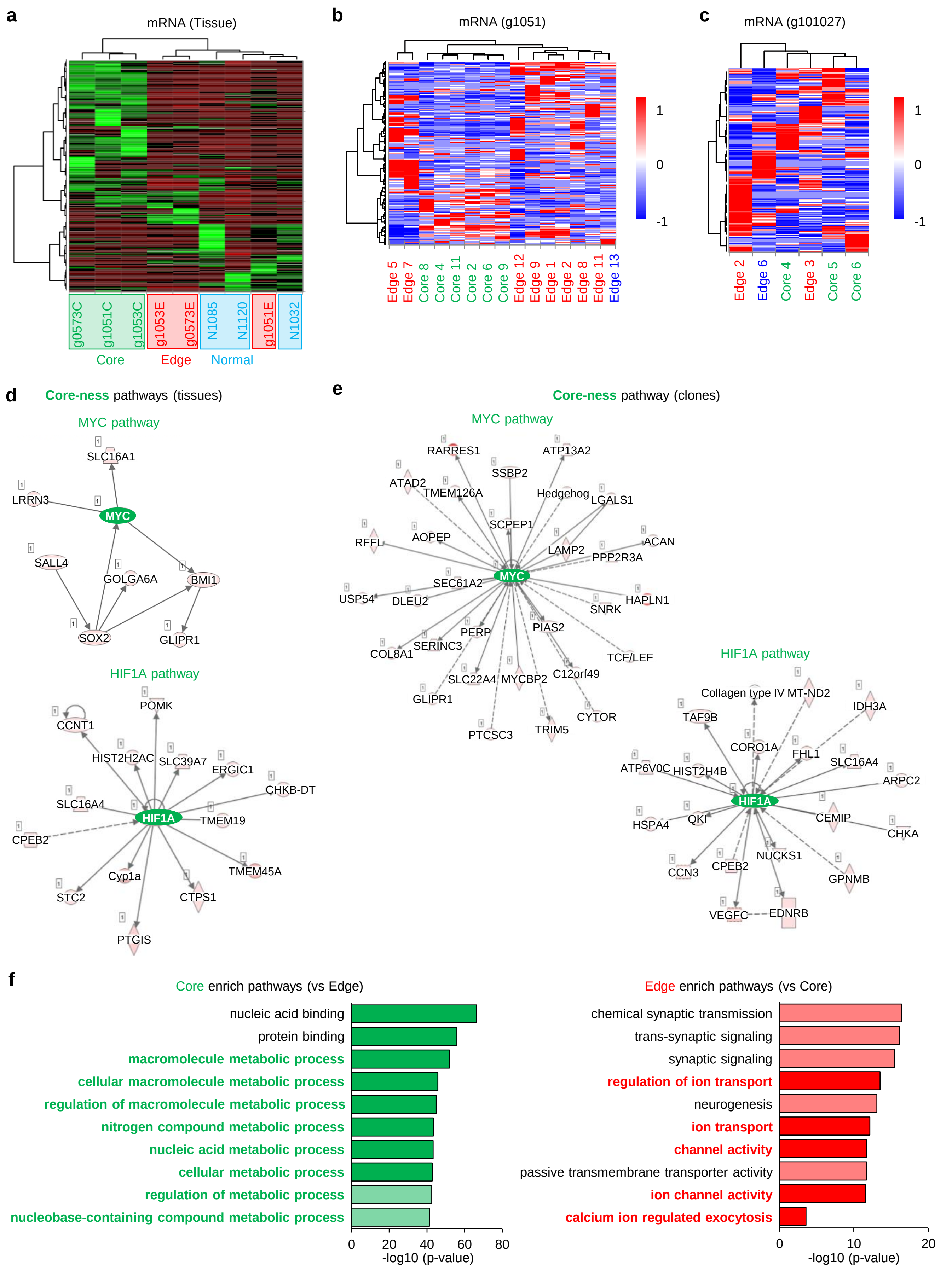

### Extended Data Figure. 3-2

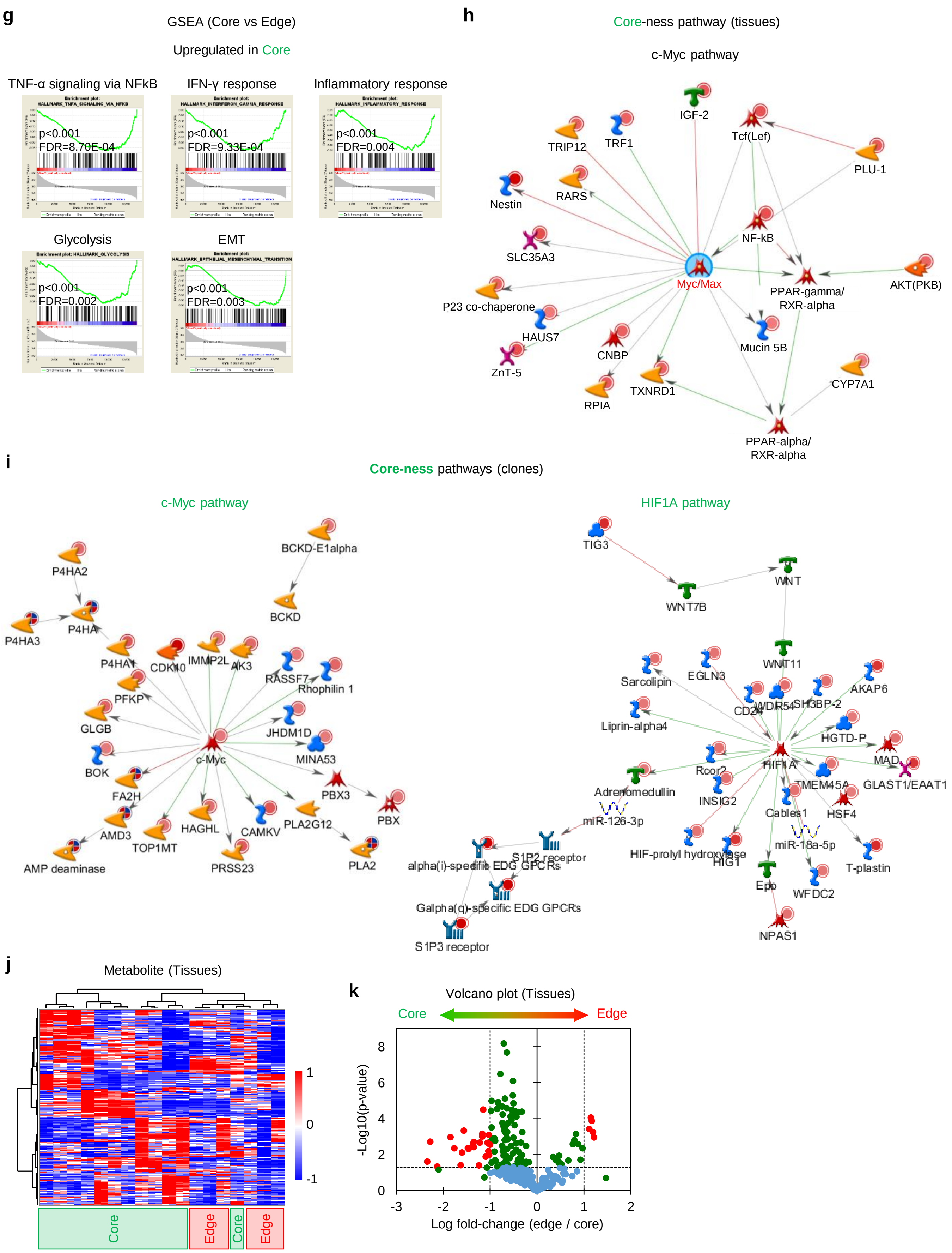

**Extended Data Figure. 3-3**

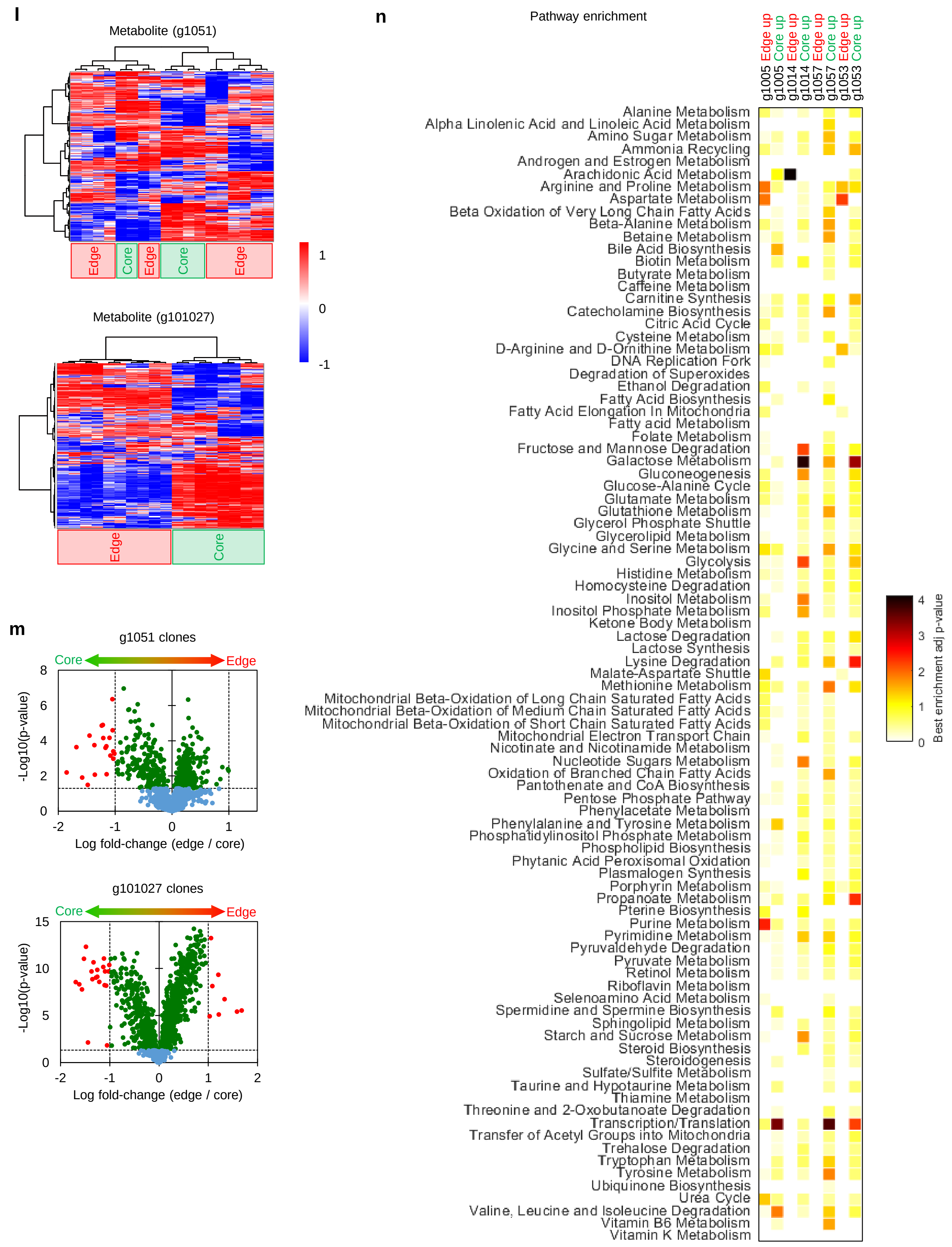

#### Extended Data Figure. 3-4

**o**

Core vs Edge (Tissues; Pt #1057)

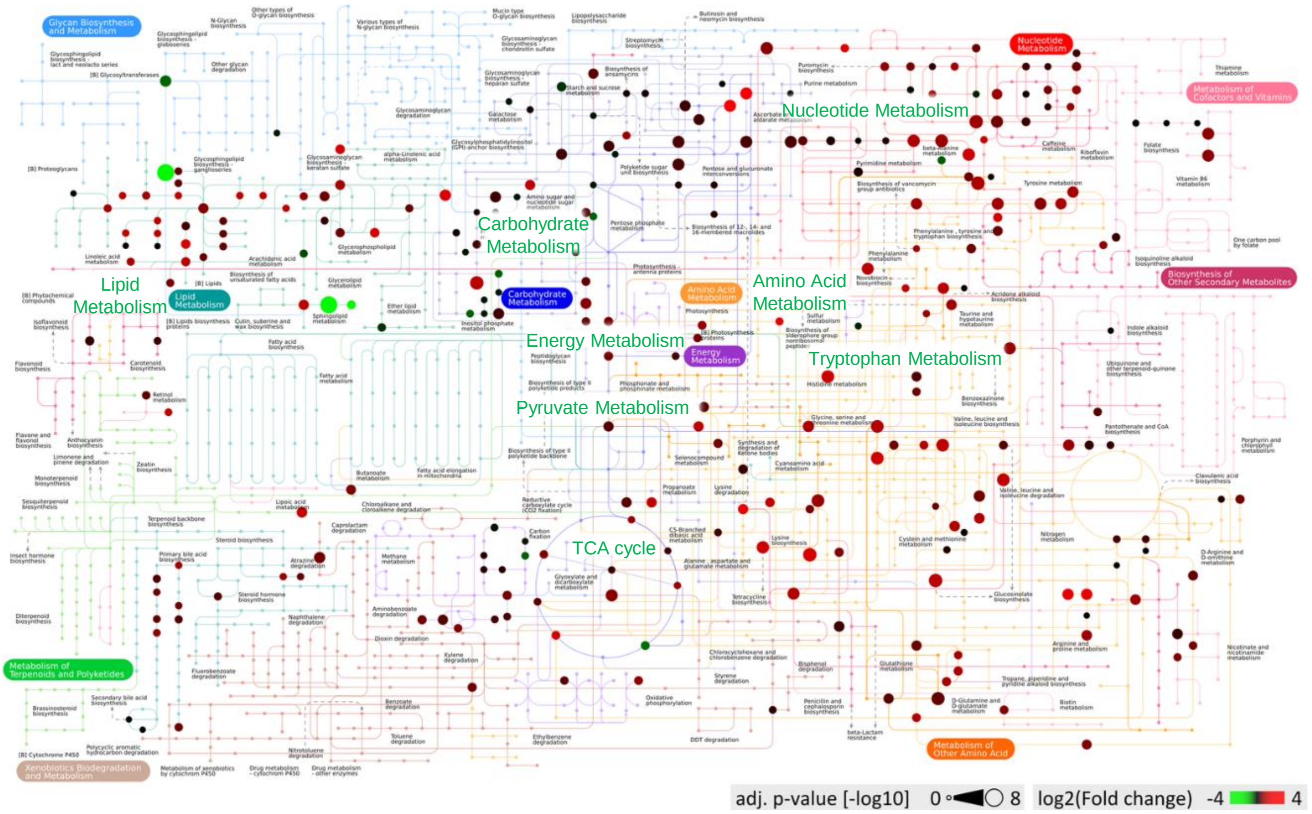

### Extended Data Figure. 3-5

**p**

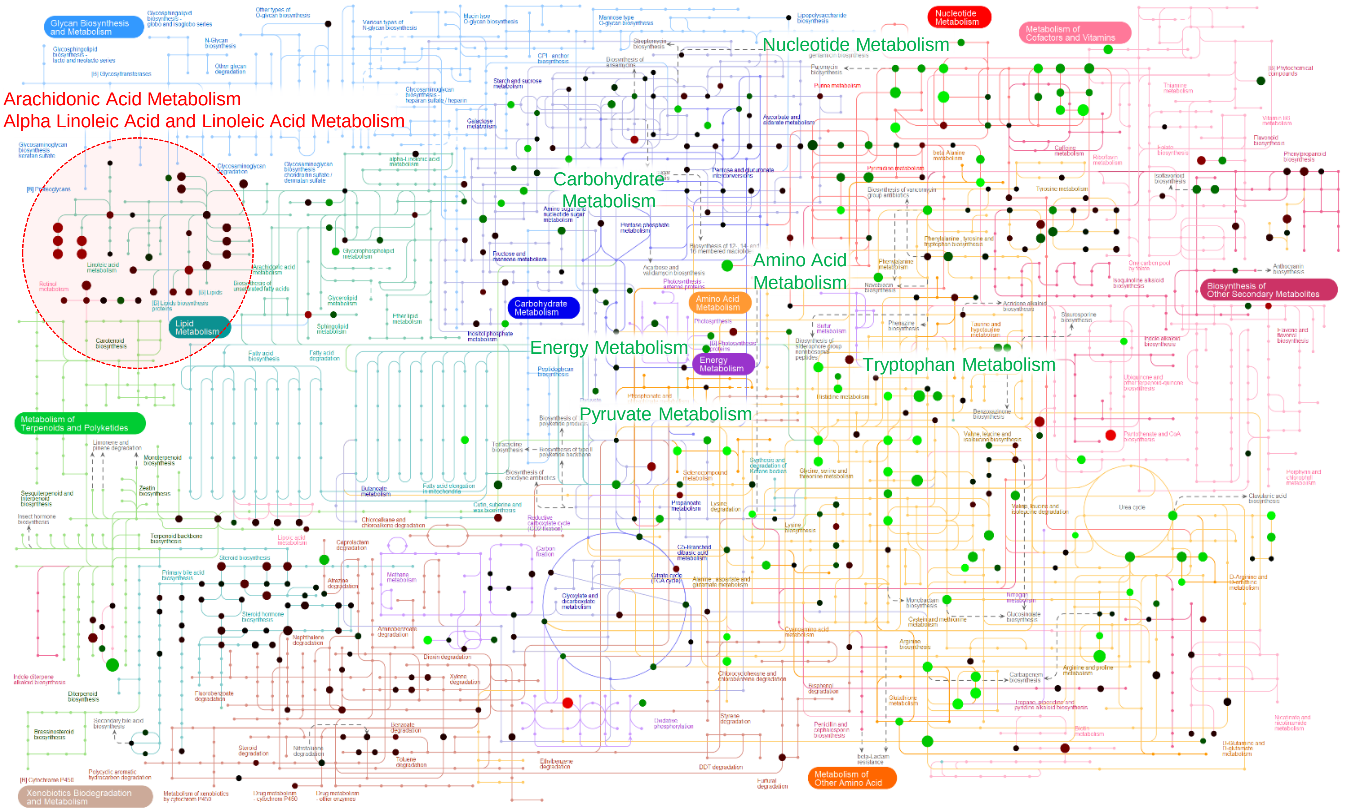


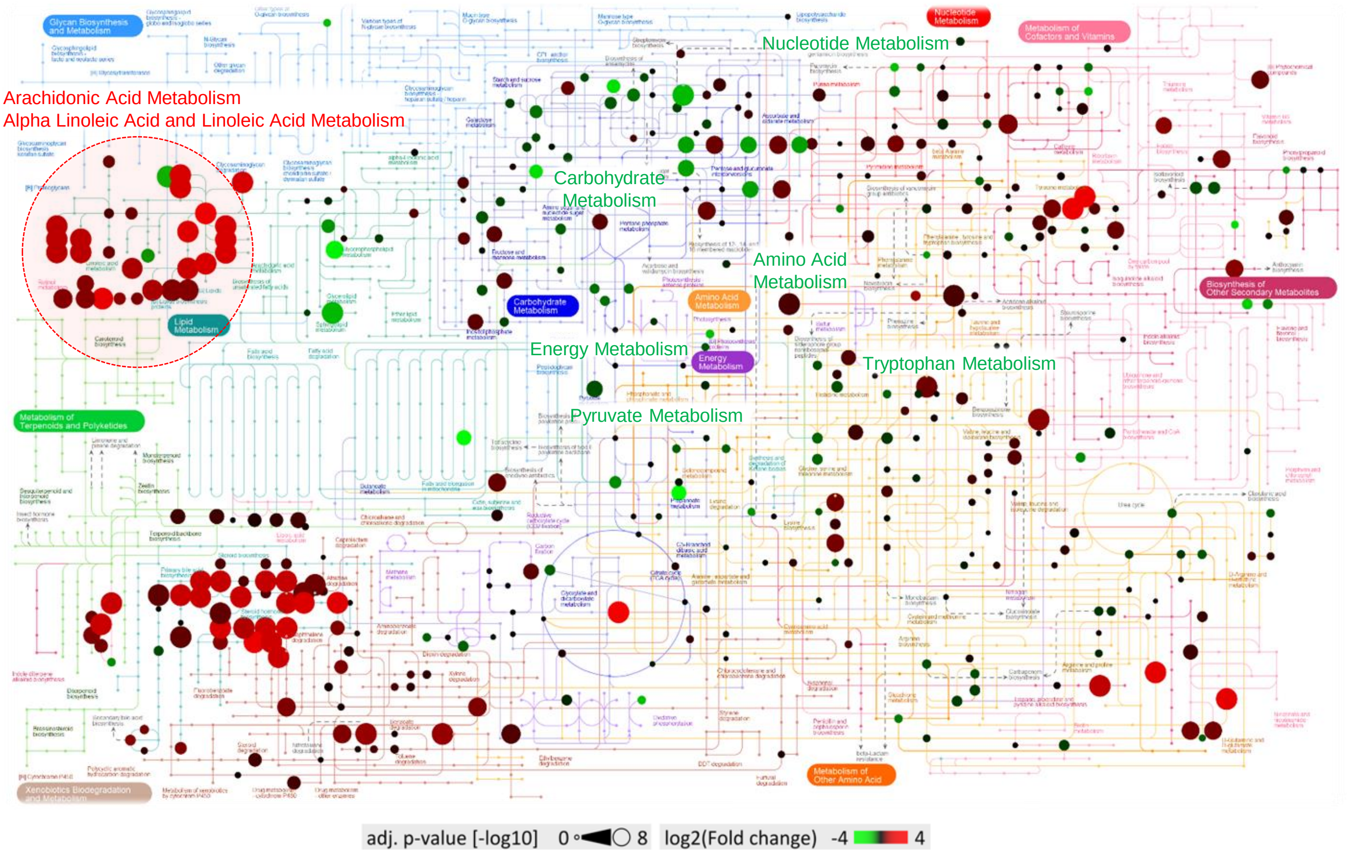

#### Extended Data Figure. 3-6

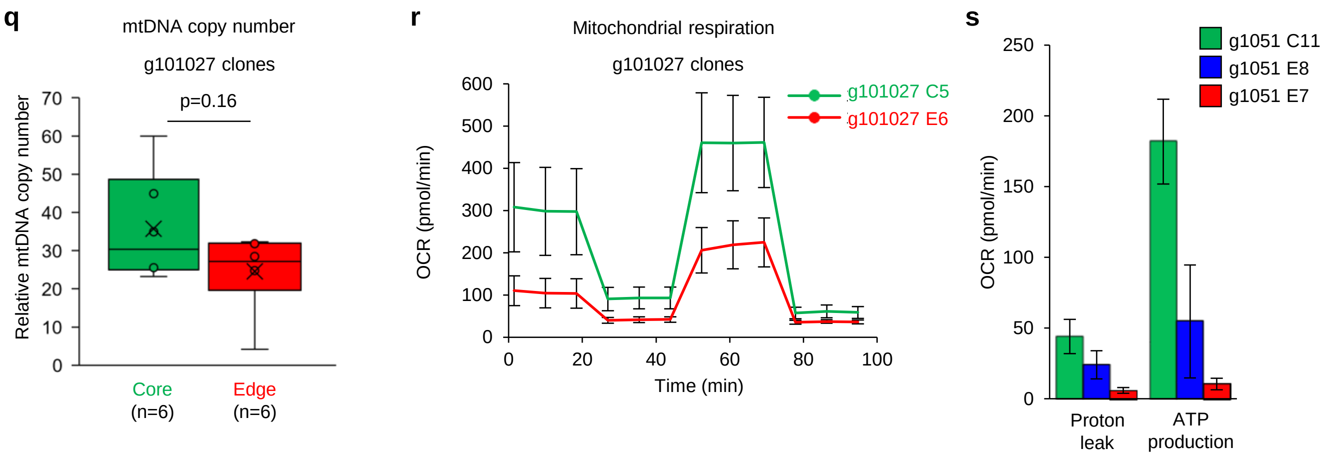

#### Extended Data Figure. 4-1

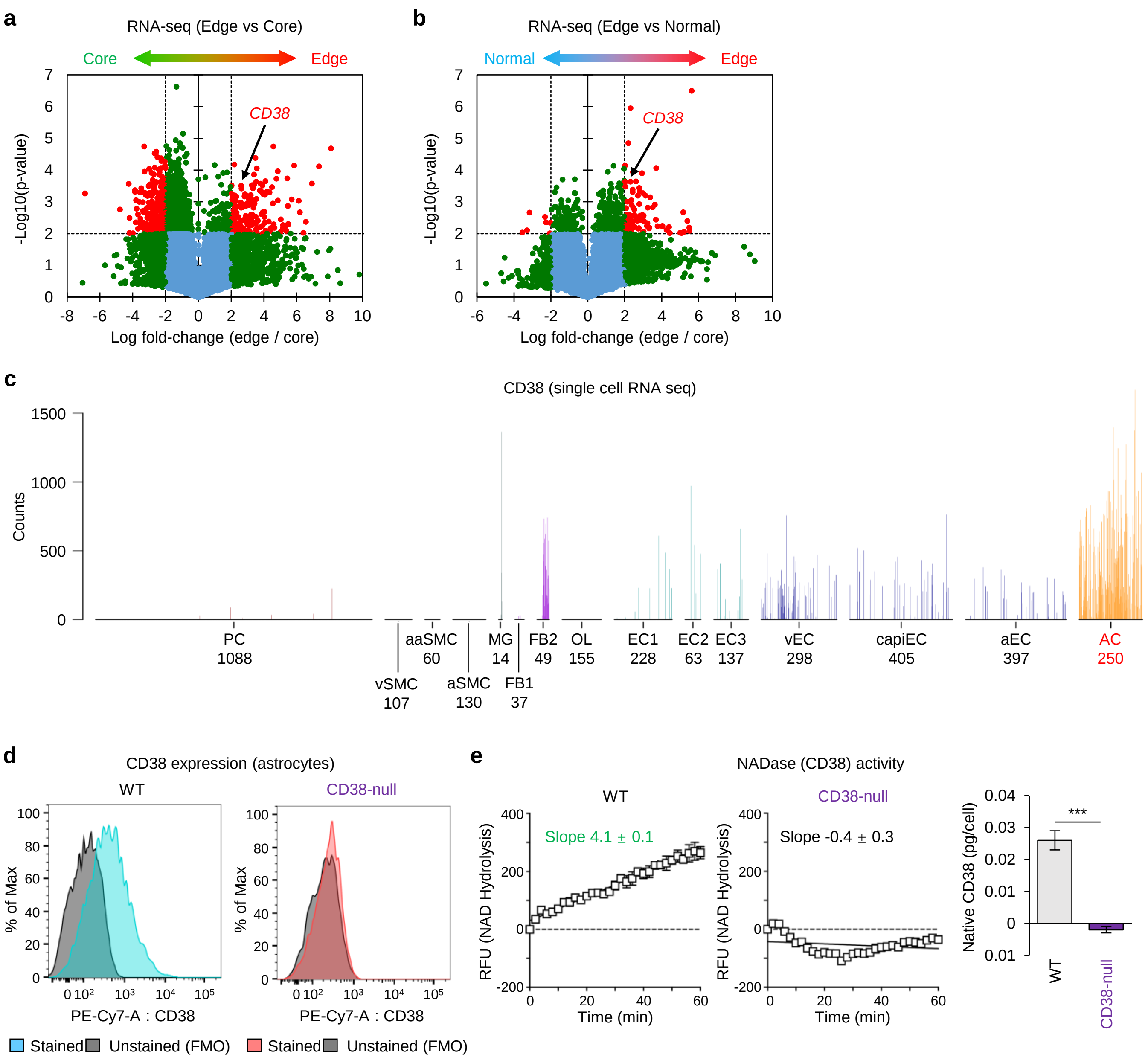

### Extended Data Figure. 4-2

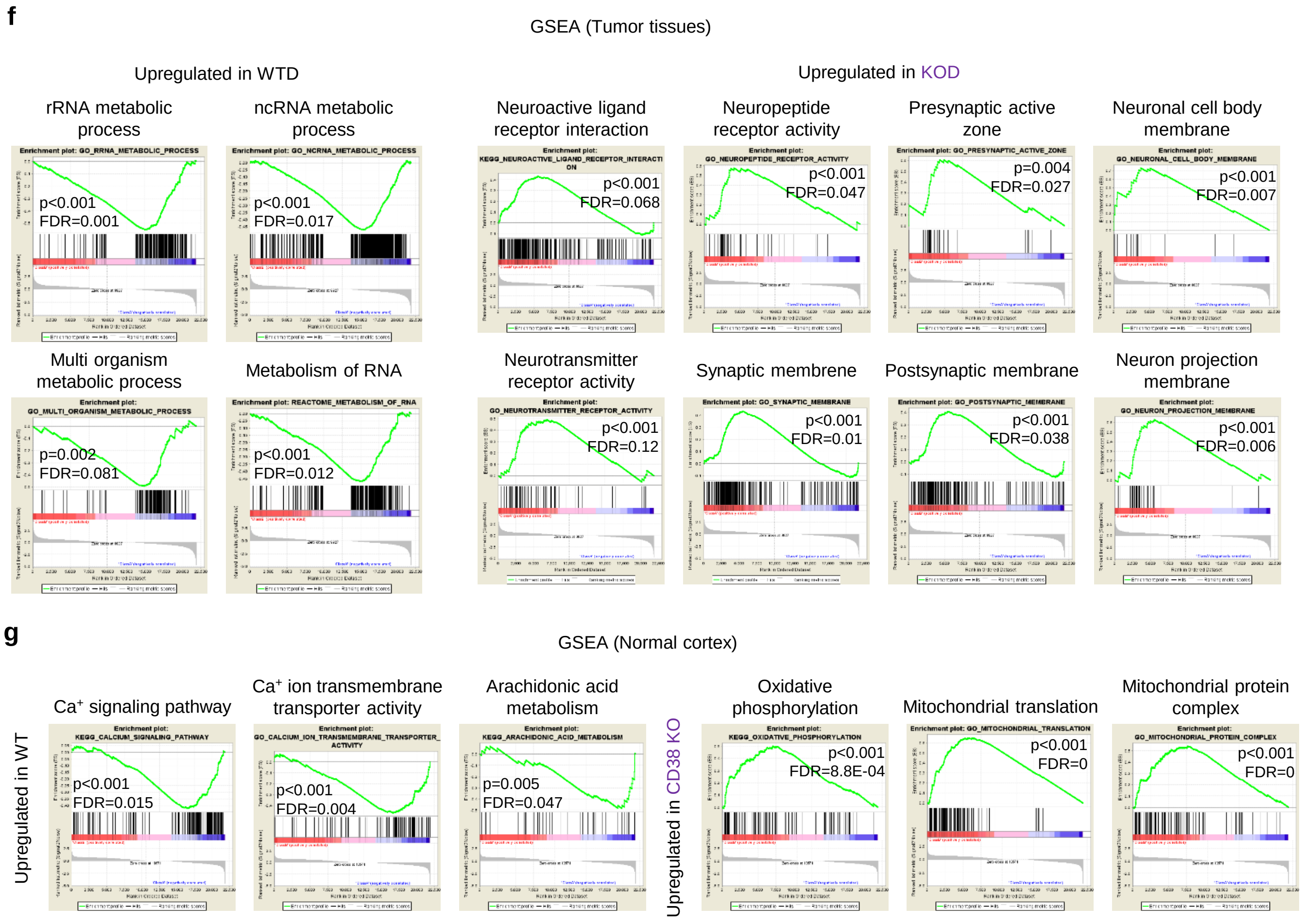

### Extended Data Figure. 6

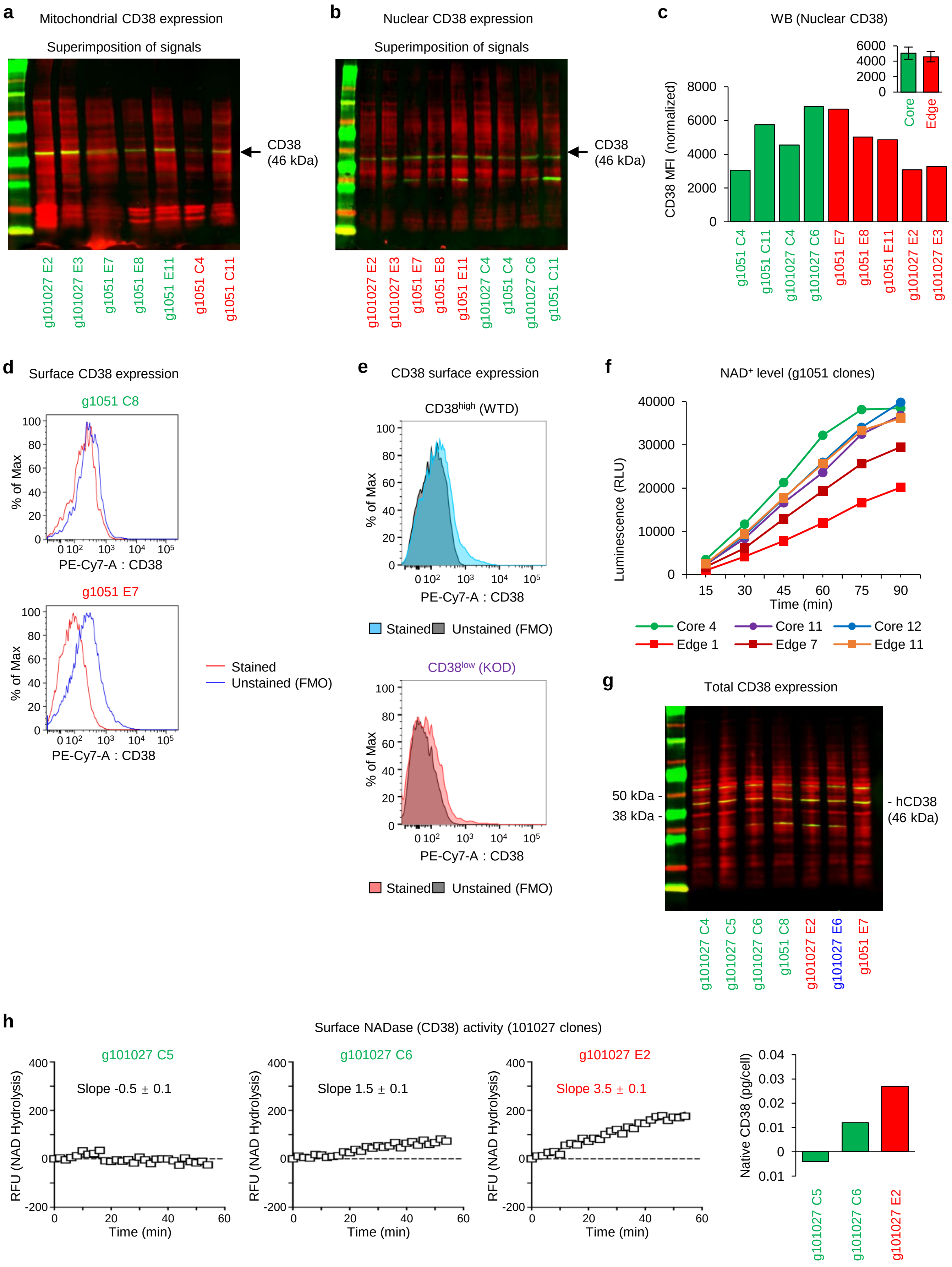

### Extended Data Figure. 7

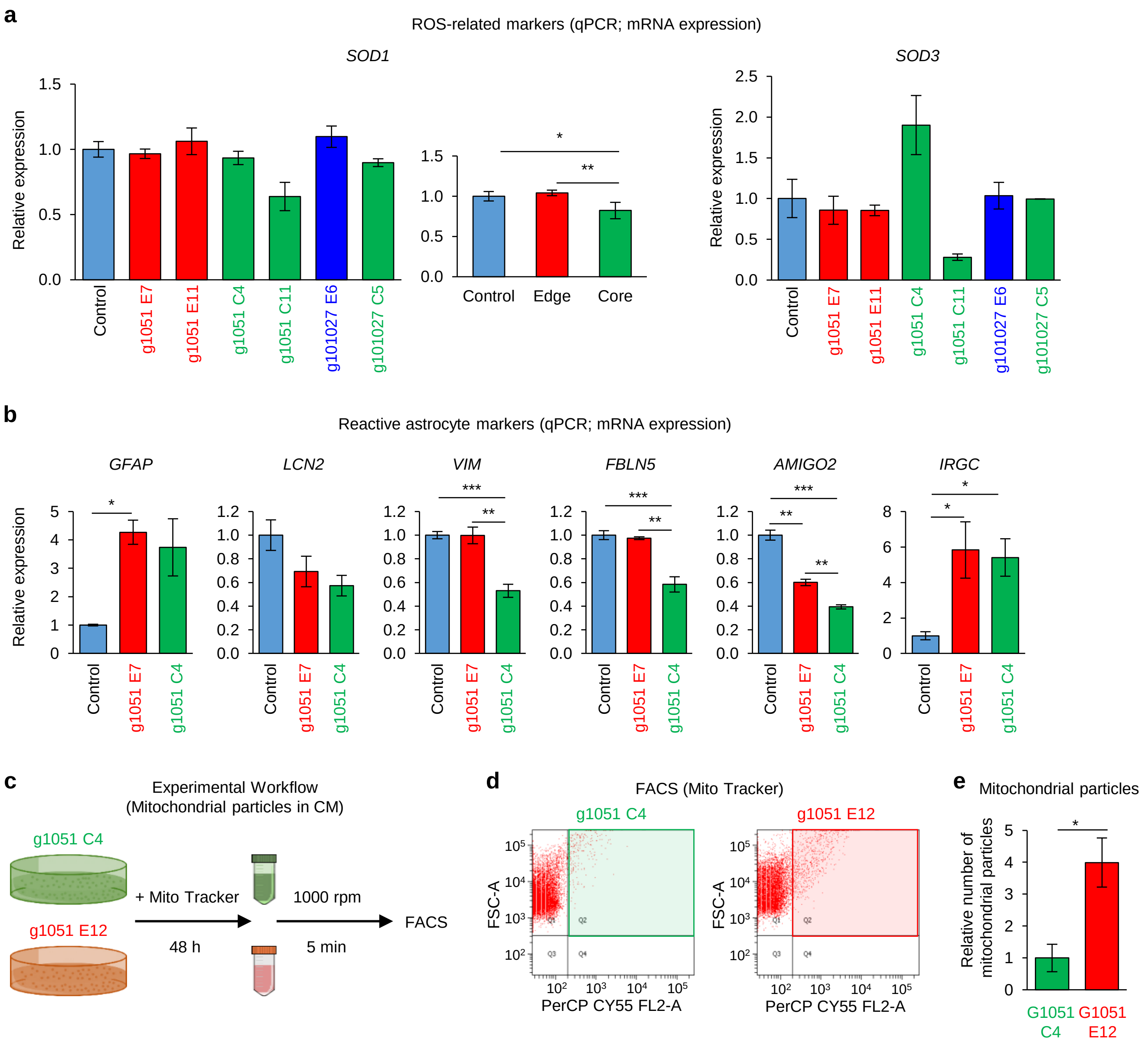

#### Extended Data Figure. 8

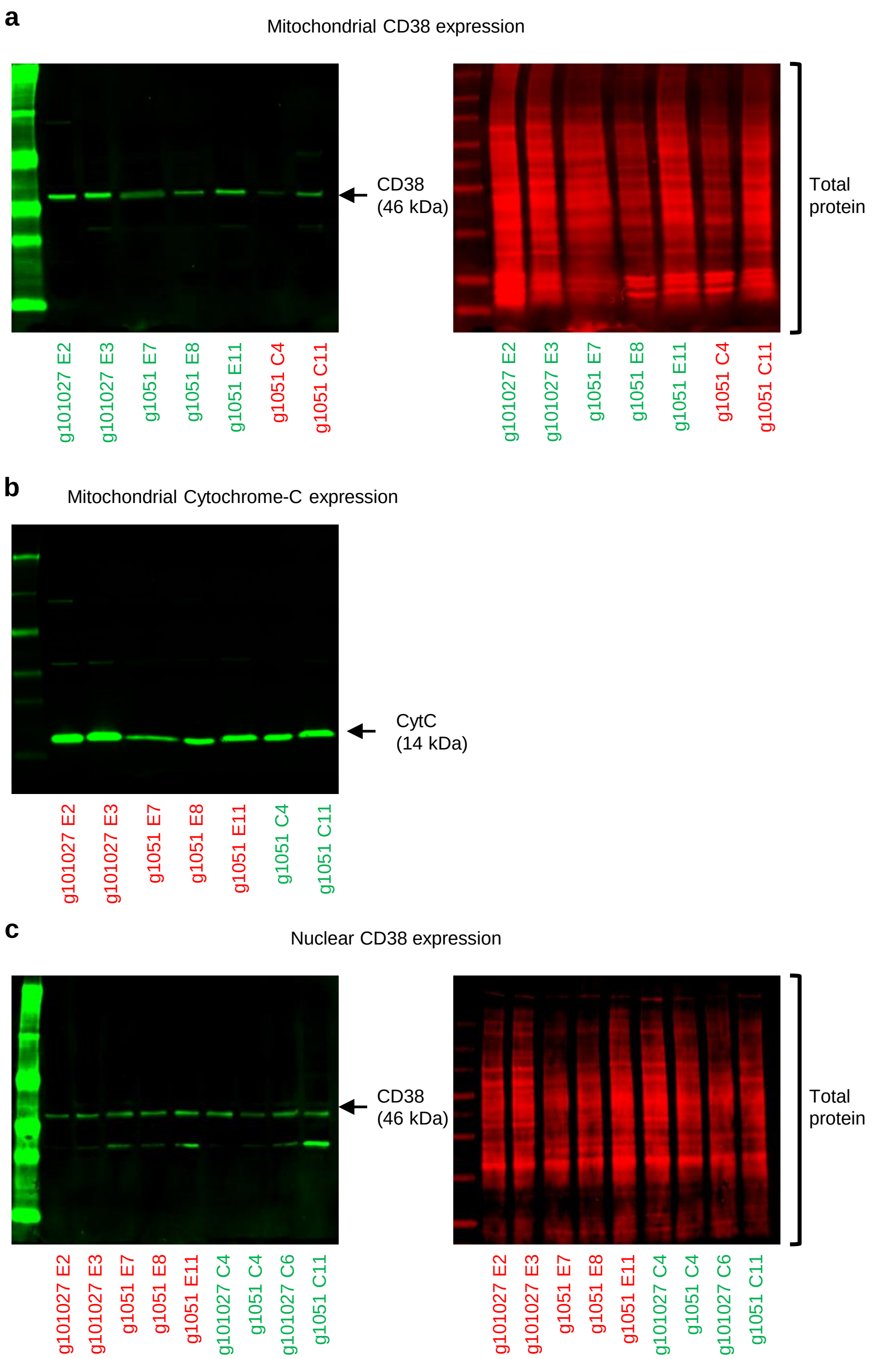
